# Targeting CD45 by gene-edited CAR-T cells for leukemia eradication and hematopoietic stem cell transplantation preconditioning

**DOI:** 10.1101/2023.10.18.562763

**Authors:** V.M. Stepanova, D.V. Volkov, D.S. Osipova, W. Wang, Y. Hou, D.E. Pershin, M.S. Fadeeva, E.A. Malahova, E.A. Kulalovskaya, L. Cuicui, Z. Mingfeng, H. Zhang, J. Xie, D. Zhang, I.Z. Mamedov, A.S. Chernov, G.B. Telegin, Y.P. Rubtsov, A.G. Gabibov, P. Wu, M.A. Maschan, A.V. Stepanov

## Abstract

Hematopoietic stem cell transplantation (HSCT) is widely used to treat patients with life-threatening hematologic and immune system disorders. The currently used nontargeted chemo-/radiotherapy conditioning regimens cause tissue injury and induce an array of immediate and delayed adverse effects, which limits the use of this potentially curative treatment. The growing demand to replace canonical conditioning regimens has led to the development of alternative approaches based on antibody‒drug conjugates, naked antibodies and CAR T cells. Here, we propose a preconditioning strategy based on targeting CD45 on hematopoietic cells with CAR45 T cells. To avoid fratricide of CD45 CAR T cells, targeted genomic disruption of the CD45 gene was performed in human CD45 CAR T cells in combination with dasatinib treatment. CD45^Δ^CAR45 T cells showed impressive activity in terms of target cell elimination *in vitro* and depletion of tumor cells *in vivo* or human hematopoietic cells in humanized immunodeficient mice engrafted with human blood-derived HSCs. CD45^Δ^CAR45 NK cells also exhibited potent killing activity against tumor cell lines and human hematopoietic cells. Therefore, fratricide-resistant CAR45 T and NK cells have the potential to provide the benefits of full myeloablative conditioning and therapy for hematologic malignancies. Thus, we provide the proof of concept for the generation and preclinical efficacy of CAR T cells directed against CD45-expressing cells.

## Key points

- Genomic disruption of CD45 prior to CAR transduction allows generation of CD45^Δ^CAR45 T cells without extensive self-antigen-driven fratricide.
- CD45^Δ^CAR45 CAR T cells can be used as a preconditioning strategy by targeting CD45 on hematopoietic cells

## Introduction

Hematopoietic stem cell transplantation (HSCT) is a curative therapeutic option for patients with hematological and immune system disorders. To achieve the successful engraftment of allogeneic HSCs, patients undergo chemo- and/or radiotherapy at doses that are highly toxic to blood and bone marrow cells but tolerated (in most patients) by other, nonhematopoietic tissues. However, graft-versus-host disease (GVHD) and nonspecific, multiorgan toxicity associated with conditioning are limiting factors for more widespread use of allogeneic HSCT^1,2^. Patients with inherited disorders such as Fanconi anemia and Nijmegen syndrome have poor tolerance to standard genotoxic preparative regimens due to hypersensitivity to DNA damage^3–5^, making HSCT risky for these patients and resulting in a particularly low success rate for patients with secondary leukemia arising from underlying DNA repair disorders.

There is considerable interest in replacing chemotherapy or radiotherapy with nongenotoxic preparative regimens. A wide range of agents, such as antibody radioconjugates^6–9^, antibody‒drug conjugates^10–13^ and naked antibodies^14–17^, have been designed to effectively deplete host stem and immune cells and enable alloengraftment. However, the eradication of highly aggressive disorders may require potent cell-based therapies. Preclinical studies have shown that myeloablation using CAR T cells specific for CD117^18,19^, CD33^20^ and CD123^21,22^ can effectively deplete HSCs and hematopoietic progenitor cells. These antigens represent a rational target for CAR T cells, as they are expressed on HSCs or hematopoietic progenitor cells, with minimal expression on nonhematopoietic tissues, and are absent on mature T lymphocytes. However, CARs targeting pan-HSCP antigens are highly desirable but not yet available. A CAR regimen that bridges HSCT preconditioning and leukemia therapy using blood tissue-specific CAR T cells could enable more comprehensive depletion of the existing hematopoietic system with a reduced risk of residual disease or relapse after transplantation.

CD45 is a cell-surface antigen that present on all immune and hematopoietic cells, including HSCs and HSPCs, except platelets and erythrocytes^23^, which makes targeting CD45 particularly suitable strategy for a conditioning regimen before HSCT. CD45 expression patterns in the bone marrow of patients with various types of acute leukemia clearly show the prevalence of CD45-positive cells^24–27^. In recent decades, CD45 has been one of the most attractive targets for nonmyeloablative regimens using antibodies, immunotoxins and ADCs^6–8,12,13,16,28–30^.

In contrast to CD117, CD33 and CD123, CD45 is among the most abundant proteins within the T-cell plasma membrane. Shared target antigen expression between CAR T cells and target cells is a significant challenge complicating CAR T-cell therapies. Since the development of CD5-specific gene-edited T cells, fratricide-resistant CAR T cells have become a rapidly growing class of leukemia immunotherapies^31^. Recent studies have demonstrated the feasibility of fratricide-resistant CARs targeting CD3^32^, CD5^31,33^ and CD7^34–38^ (CD5/CD7 bispecific^35^) and CD38^39,40^. Some of these approaches are being tested in early-phase clinical trials, with encouraging results (#NCT04004637, #ChiCTR2000034762, #NCT03690011; clinicaltrials.gov).

Here, we describe the generation of CAR T cells directed against human CD45 as a proof-of-concept for leukemia therapy and nongenotoxic conditioning prior to HSCT. To improve the translational potential of this approach, the CAR45 construct was designed to consist of the anti-CD45 antibody BC8, which has been extensively tested in clinical trials (#NCT00002554, #NCT00003868, #NCT00003870, #NCT00005940, #NCT00008177, #NCT00988715, #NCT01300572, #NCT01503242, #NCT01921387; clinicaltrials.gov) fused with a 3^rd^-generation CAR backbone.

To overcome CD45-dependent fratricide during the CAR45 T-cell manufacturing process and after infusion, we performed targeted disruption of the CD45 gene using CRISPR/Cas9. We showed that our gene-editing and CAR transduction protocol consistently yields a highly enriched population of CD45-negative CAR45 T and NK cells (CD45^Δ^CAR45). The CD45^Δ^CAR45 T cells demonstrate robust antitumor activity against hematologic malignancies and normal human hematopoietic cells *in vitro* and *in vivo*. If successful, CD45^Δ^CAR45 T cells could serve as an alternative nongenotoxic tool to eradicate both normal and pathologic hematopoietic cells, ensuring engraftment and clearance of the leukemia burden while sparing nonhematopoietic tissue, mature platelets, and red blood cells.

## Methods

### Flow cytometry analysis

A total of 3×10^5^ cells were centrifuged (300×g; 5 min; RT) and resuspended in 100 μL of PBS. Surface antigens were stained with fluorescently labeled antibodies (**Supplementary Table 1**) for 30 min at 4 °C. The cells were washed once with 1 mL of cold PBS (300×g, 5 min, RT) and resuspended in 110 μL of cold PBS. Data were acquired using an ACEA NovoCyte 2060 flow cytometer (Agilent, USA). ACEA NovoExpress (Agilent, USA) was used for data analysis. For FACS, stained cells were acquired on an SH800 Cell Sorter (Sony, USA).

### Cells and culture conditions

The HEK293T lentivirus packaging (Clontech, USA) and 293Vec-RD114 retrovirus packaging cell lines (BioVec Pharma, Canada) were cultured in DMEM (Gibco, USA) supplemented with 10% FBS (Gibco, USA), 100 U/mL penicillin (Gibco, USA), 100 μg/mL streptomycin (Gibco, USA), and 2 mM GlutaMAX (Gibco). The Jeko-1, Nalm-6, Jurkat, Raji, THP-1, K562 and Ramos cell lines were cultured in RPMI 1640 (Gibco, USA) supplemented with 10% FBS (Gibco, USA), 100 U/mL penicillin (Gibco, USA), 100 μg/mL streptomycin (Gibco, USA), and 2 mM GlutaMAX (Gibco). The lentivirus packaging HEK293T cell line was purchased from Clontech, USA. The Jeko-1, Nalm-6, Raji, THP-1, K562, Ramos and Jurkat cell lines were purchased from ATCC, USA. The Jurkat NFAT Lucia reporter cell line was purchased from Invivogen, USA. Irradiated K562 feeder cells with 4-1BB and IL-21 surface expression were obtained as described previously^41^.

T cells from healthy donors were activated and cultured in TexMacs medium (Miltenyi, USA) supplemented with 12.5 ng/mL IL-7 (Miltenyi, USA) and 12.5 ng/mL IL-15 (Miltenyi, USA) or RPMI 1640 (Gibco, USA) medium supplemented with 100 U/mL IL-2 (Sci-Store, Russia). NK cells from healthy donors were cultured in TexMacs medium (Miltenyi, USA) supplemented with 10 U/mL IL-2 (Sci-Store, Russia). All cytotoxicity assays were performed in TexMacs medium with IL-7 and IL-15 or RPMI 1640 medium with IL-2 (Sci-Store, Russia). The Jeko-1, Nalm-6, Raji, THP-1, K562, Ramos and Jurkat cell lines were modified by lentiviral transduction to induce stable firefly luciferase (Ffluc) or green fluorescent protein (GFP) expression. For the Incucyte assay, THP-1 or Jeko-1 cells were transduced with IncuCyte NucLight Red Lentivirus Reagent (EF-1 Alpha, Puro) (Sartorius, USA). All cell lines were repeatedly tested for the presence of mycoplasma contamination with a MycoReport Mycoplasma Detection Kit (Evrogen, Russia).

### Isolation of primary human peripheral blood mononuclear cells

Human peripheral blood mononuclear cells (PBMCs) were isolated from the blood of healthy donors by gradient density centrifugation on Ficoll-Paque (GE Healthcare, USA) according to a standard protocol approved by the local ethics committee of the Dmitry Rogachev National Medical Research Center of Pediatric Hematology, Oncology, and Immunology. All participants provided informed consent. T cells were isolated from human PBMCs with an Untouched Human T-cell Isolation Kit (Invitrogen, USA) and activated with Dynabeads Human T Activator CD3/CD28 (Gibco, USA) at a 1:1 ratio for 24 hours. NK cells were isolated from PBMCs with an NK Cell Isolation Kit (Miltenyi, USA). NK cells were mixed with irradiated K562 feeder cells at a 1:10 ratio for 5 days, and on day 5, transduction was performed.

### Chimeric antigen receptors

The nucleotide sequence of CAR45 consists of the following parts: anti-CD45 scFv (from CD45 antibody clone BC8) ^42^, the human IgG4 hinge region, CD28 transmembrane domain, CD28 and OX-40 costimulatory domains, and the CD3ζ signaling domain. A synthetic anti-CD45 scFv gene (GeneCust, France) was cloned and inserted into the pLV2 lentiviral vector (Clontech, USA) and pMM25b retroviral vector (a gift from Dr. M. Mamonkin). The CAR19 nucleotide sequences were obtained as described previously^43^ and cloned and inserted into the pLV2 and pMM25b vectors. All sequences were validated by Sanger sequencing (Evrogen, Russia).

### Transduction of T cells and NK cells with pseudoviral particles

Lentiviral particles containing CAR45 and CAR19 were produced by polyethylenimine-mediated cotransfection (Sigma, USA) of HEK293T cells with the corresponding lentiviral CAR plasmids and the packaging plasmids pCMV VSV-G, pRSV-REV, and pMDLg-pRRE (a gift from Dr. I. Verma). Supernatants were collected at 48 hours post transfection and purified by 2 rounds of centrifugation (1st - 300×g, 5 min, RT; 2nd - 4500×g, 5 min, RT). On the 2nd day after isolation, 1×10^6^/mL activated T cells were resuspended in TexMacs media and mixed with lentiviral supernatants and 13 μg/mL Polybrene (Sigma, USA). Dasatinib (50 nM, Stemcell, Canada) was added prior to transduction. The cells were centrifuged at 1200×g for 90 min at 32 °C and incubated at 37 °C with 5% CO_2_ for 18 hours. The culture medium was changed on the day after transduction and every 3-4 days to maintain the cells at a density of 2.0×10^6^ cells/mL.

Retroviral particles containing CAR45 or CAR19 were produced by polyethylenimine-mediated transfection of 293Vec-RD114 cells. Supernatants containing the virus were collected at 48 hours post transfection and purified by 2 rounds of centrifugation (1st - 300×g, 5 min, RT; 2nd - 4500×g, 5 min, RT). A total of 0.5×10^6^ NK cells were resuspended in TexMacs medium with 100 nM dasatinib on the 5th day after isolation. Retroviral supernatants were mixed with 10 μg/mL Vectofusin-1 (Miltenyi, USA) and added to NK cell suspensions. The cells were centrifuged at 400×g for 2 hours at 37 °C and incubated at 37 °C with 5% CO_2_ for 48 hours. The culture medium was changed on the 2nd day after transduction and every 3-4 days to maintain cells at a density of 2.0×10^6^ cells/mL.

### Target cell modification

To obtain reporter cells, Jeko-1, Nalm-6, Nalm-6 CD19 ^Δ^, Raji, THP-1, K562, Ramos and Jurkat cells were transduced to express the fluorescent protein GFP and the reporter protein Firefly Luciferase simultaneously. The reporter cell lines were stained with anti-human CD45 and anti-human CD19 antibodies and analyzed by flow cytometry on an ACEA NovoCyte 2060 (Agilent, USA). ACEA NovoExpress (Agilent, USA) was used for data analysis.

### Gene knockout in cells

The procedure was performed by electroporation with a 4D-Nucleofector device (Lonza, Switzerland) using the CRISPR/Cas9 system and the P3 Primary Cell 4D-Nucleofector X Kit L (Lonza, Switzerland). Oligonucleotides for gRNA were selected by Сrispor (Tefor Infrastructure, France) algorithms and synthesized by Evrogen, Russia (**Supplementary Figure 1a**). Three gRNAs specific to the human CD45 gene (GGATTTGTGGCTTAAACTCT, GGGTTTAAGCCACAAATACA and GGCACACTTATACTCATGTT) were evaluated for gene editing efficiency in combination and/or individually. Two gRNAs (GGCTCATGAGCTTCCCGGAA and GGGCGGGGACTCCCGAGACC) were utilized for CD19 gene knockout as described previously^44^. Short nucleotide sequences were used for transcription according to the HiScribe Quick T7 High Yield RNA Synthesis Kit (NEB, UK) protocol. The resulting gRNAs were purified using the AMPure XP kit and treated with DNase I (NEB, UK). Concentration was evaluated on a Qubit system (Thermo Fisher Scientific, USA). For electroporation, 2 μg of gRNA was mixed with 3.3 μL of Spy Cas9 NLS (NEB, UK) and incubated for 30 min at RT. For CD45 knockout, T and CAR T cells (on the 3rd day after isolation and the 2nd day after transduction) or NK cells and CAR NK cells (on the 7th day after isolation and the 2nd day after transduction) were washed twice with PBS (300×g, 5 min, RT), resuspended in 100 μL of electroporation buffer and mixed with the gRNA + Cas9 mix according to the P3 protocol for the Primary Cell 4D-Nucleofector XL Kit. For CD19 knockout, Nalm-6 cells were prepared similarly. Electroporation was performed using the DN-100 program for NK cells and the EH100 program for T cells and Nalm-6 cells. The survival and proliferation rates were evaluated by trypan blue 0.04% (Bio-Rad, USA) staining every 2 days using a TC20 cell counter (Bio-Rad, USA). Briefly, cells were resuspended in culture medium, and 10 μL of the suspension was mixed with 10 μL of trypan blue. Ten microliters of the mixture was injected into a counting slide (Bio-Rad, USA) and analyzed on a cell counter. The knockout efficiency was assessed on the 5th day after electroporation by flow cytometry on an ACEA NovoCyte 2060 (Agilent, USA). ACEA NovoExpress (Agilent, USA) was used for data analysis. For Nalm-6 cells, the knockout population was enriched by FACS on an SH800S Cell Sorter (Sony, USA).

### Off-target analysis for CD45 gRNAs

A total of 183 potential off-target sites were identified with the use of the CRISPOR software tool for off-target prediction. We selected all predicted off-target sites with a CFD (cutting frequency determination) score ≥ 0.21 (N = 20) and the off-target sites predicted within exons independently of the CFD (N=5). For each of the selected loci, we designed a pair of primers using Primer-BLAST. A pair of primers flanking the guide RNA target in PTPRC was added as a positive control. These primers were pooled and used for multiplex PCR with genomic DNA isolated from cells electroporated with Cas9/guide RNA RNP or with pure Cas9 without guide RNA as a negative control. Cells were collected at 4, 8, and 16 hours after electroporation. Six multiplex amplicons were ligated to Illumina TruSeq adapters using the NEBNext Ultra II DNA Library Prep Kit for Illumina (NEB), pooled, and sequenced on an Illumina NextSeq 550 (paired-end 100+100), resulting in 2.1-2.8 million raw sequencing reads per library. The reads were preprocessed with TrimGalore v0.6.4 and mapped to the human genome (Hg38) using Bowtie 2 v2.3.5.1. Metrics for target regions were collected with GATK v4.1.4.1 CollectTargetedPcrMetrics. The percentage of insertions and deletions was calculated in R with the GenomicAlignments package for each amplicon separately in cells treated with Cas9 with and without guide RNA at different timepoints.

### Sanger sequencing

gDNA for T and CD45^Δ^ T cells was obtained with an ExtractDNA Blood & Cells Kit (Evrogen, Russia) on the 2^nd^ day after knockout. The BigDye® Direct Sanger Sequencing Kit (Thermo Fischer Scientific, USA) and GCAAAGAGGACCCTTACAGTAT and AGATACAGATAGACATTGCTTTC primers were utilized for the subsequent analysis according to the manufacturer’s instructions. Sequencing reactions were analyzed on a 3500 Genetic Analyzer (Thermo Fischer Scientific, USA).

### Western blotting

Cell pellets of 1×10^5^ T or CD45^Δ^ T cells were lysed with RIPA lysis buffer (50 mM Tris HCl, 150 mM NaCl, 1.0% (v/v) NP-40, 0.5% (w/v) sodium deoxycholate, 1.0 mM EDTA, 0.1% (w/v) SDS and 0.01% (w/v) sodium azide at a pH of 7.4). Cell lysates were mixed with 2× sample buffer (125 mM Tris·HCl pH 6.8, 4% (w/v) SDS, 20% (v/v) glycerol, 100 mM DTT, 0.01% bromophenol blue) and heated at 90 °C for 10 min. Samples were separated by SDS‒PAGE on a 12.5% polyacrylamide gel, transferred to a nitrocellulose membrane (Bio-Rad, USA) and incubated with a recombinant anti-human CD45 primary antibody and an HRP-conjugated goat anti-rabbit IgG secondary antibody according to the manufacturers’ protocols. An HRP-conjugated anti-human GAPDH antibody was used as a positive control. The presence of CD45 and GAPDH was assessed on a VersaDoc 4000 MP Imaging System (Bio-Rad, USA) using an Enhanced Chemiluminescence kit (Bio-Rad, USA).

### Confocal microscopy

Jurkat T cells or CD45^Δ^ Jurkat T cells were plated on poly-L-lysine-covered coverslips and centrifuged (100×g, 10 min, RT). The attached cells were fixed in 4% formalin for 1 hour at RT and stained with anti-human CD45 antibodies against extracellular and intracellular CD45. Confocal images were captured using an Axio Observer Z1 microscope (Carl Zeiss AG, Germany) with a Yokogawa spinning disc confocal device CSU-X1 (Yokogawa Corporation of America, USA).

### CAR T-cell expansion assay

A total of 1×10^5^ CD45^Δ^ CAR45 T cells or CD45^Δ^ CAR19 T cells were plated in 24-well plates in TexMacs medium supplemented with IL-7/IL-15 and with or without 50 nM dasatinib (Stemcell, Canada). Cell number and viability were assessed every 2 days for 2 weeks by flow cytometry on an ACEA NovoCyte 2060 (Agilent, USA). ACEA NovoExpress (Agilent, USA) was used for data analysis.

### Detection of proinflammatory cytokines

A total of 5×10^4^ Ramos cells (target cells) were mixed with 2.5×10^5^ T or CD45^Δ^ T cells (effector cells) on the 8th-16th days after knockout of CD45 (E:T ratio 5:1) in TexMacs medium supplemented with IL-7, IL-15, and 1 nM blinatumomab (Amgen, USA) in the absence of IL-2. Wells containing only effector cells were used as negative controls. Cells were incubated at 37 °C with 5% CO_2_ for 24 hours. The supernatant was separated from the cells by centrifugation (4 °C, 300×g, 5 min) and stored at −20 °C before use. The concentrations of the cytokines IL-2 and IFN-γ were measured using ELISA-based kits (Vector-best, Russia) according to the manufacturer’s instructions. Data were analyzed with a Varioskan Flash instrument (Thermo Fischer Scientific, USA) using SkanIt RE for Varioskan Flash 2.4.5 (Thermo Fischer Scientific, USA).

A total of 5×10^3^, 1×10^4^, 2.5×10^4^ or 5×10^4^ THP-1 or Jeko-1 cells (target cells) were mixed with 5×10^4^ CD45^Δ^ CAR45 T cells, CD45^Δ^ CAR19 T cells, or CD45^Δ^ T cells (effector cells) on the 10th day after CD45 knockout (E:T ratios 10:1, 5:1, 2:1, 1:1) in RPMI 1640 medium. Wells containing only effector cells were used as a negative control. The cells were incubated at 37 °C with 5% CO_2_ for 24 hours. The supernatant was separated from the cells by centrifugation (4 °C, 300×g, 5 min) and stored at −20 °C before use. The concentrations of the cytokines IL-2 and IFN-γ were measured using ELISA-based kits (Vector-best, Russia) according to the manufacturer’s instructions. Data were analyzed with a Varioskan Flash instrument (Thermo Fischer Scientific, USA) using SkanIt RE for Varioskan Flash 2.4.5 (Thermo Fischer Scientific, USA).

### T and CD45^Δ^ T-cell cytotoxicity assay

A total of 5×10^4^ Ramos cells expressing GFP (target cells) were mixed with 5×10^4^ T or CD45^Δ^ T cells (effector cells) (E:T ratio 1:1) in TexMacs medium supplemented with IL-7, IL-15 and 1 nM blinatumomab. Wells with target cells only were used as a negative control. The cells were incubated at 37 °C with 5% CO_2_ for 24 hours and were washed twice with PBS (Gibco, USA) (RT, 300×g, 5 min). The results were evaluated by flow cytometry on an ACEA NovoCyte 2060 (Agilent, USA). ACEA NovoExpress (Agilent, USA) was used for data analysis.

### Cytotoxicity assays of CAR T cells and CAR NK cells

#### Tumor cells

A total of 1×10^4^, 2×10^4^ or 5×10^4^ CD19 ^Δ^ Nalm-6, Nalm-6, Raji, Jeko-1, Jurkat, THP-1, or Ramos cells (target cells) were mixed with 5×10^4^ CD45^Δ^ CAR45 T cells, CAR45 T cells, CD45^Δ^ CAR19 T cells, CAR19 T cells, or T cells (effector cells) on the 8th-16th days after knockout of CD45 (E:T ratios were 5:1, 2.5:1, 1:1). Wells containing only target cells were used as a negative control for cytotoxicity. Wells containing only effector cells were used as a positive control for viability. The cells were incubated at 37 °C with 5% CO_2_ for 24 hours and washed twice with PBS (Gibco, USA) (300×g, 5 min, RT). The results were evaluated by flow cytometry on an ACEA NovoCyte 2060 (Agilent, USA). ACEA NovoExpress (Agilent, USA) was used for data analysis.

#### Incucyte killing assay

For the Incucyte killing assay, 3×10^4^ red fluorescent THP-1 or Jeko-1 cells (target cells) were mixed with 9×10^4^ CD45^Δ^ CAR45 T cells, CAR45 T cells, CD45^Δ^ CAR19 T cells, CAR19 T cells, or T cells (effector cells) on the 8th-16th days after knockout (E:T ratio was 3:1) in TexMacs medium (Miltenyi, USA) and incubated at 37 °C with 5% CO_2_ for 48 hours. Wells containing only target cells were used as negative controls. Changes in fluorescence were monitored every 2 hours using the IncuCyte Live-Cell Analysis System (Essen BioScience, USA). To determine the cytolytic activity of effector cells, the area under the curve (AUC) was calculated.

#### T cells and PBMCs

A total of 3×10^4^ T cells or PBMCPBMCs (target cells) were mixed with CD45^Δ^ CAR19 T cells, CD45^Δ^ CAR45 T cells or CD45^Δ^ T cells (E:T ratio 1:5, 1:2, 1:1, 5:1) or CD45^Δ^ CAR45 NK cells, CAR45 NK cells or NK cells (effector cells) on the 8th-16th days after CD45 knockout (E:T ratios – 5:1, 1:1, 1:2). Prior to the experiment, target T cells and PBMCs were stained with 5 μM CFSE (Invitrogen, USA) according to the manufacturer’s protocol. Wells containing only target or only effector cells were used as negative controls. For the CAR T-cell experiment, the cells were incubated at 37 °C with 5% CO_2_ for 24 hours. For the CAR NK cell experiment, the cells were incubated at 37 °C with 5% CO_2_ for 4 and 24 hours. One hundred microliters of cell suspension from each well was stained with anti-human CD45 (anti-human CD3 for NK cell experiments) according to the manufacturer’s instructions and washed twice with PBS (Gibco, USA) (300×g, 5 min, 4 °C). The results were evaluated by flow cytometry using ACEA NovoCyte 2060 (Agilent, USA). ACEA NovoExpress (Agilent, USA) was used for data analysis.

#### PBMCs

PBMCs (3×10^5^, target cells) were mixed with CD45^Δ^ CAR45 T cells or CD45^Δ^ T cells (E:T ratio 1:3) on the 8th-16th days after CD45 knockout. Wells containing only PBMCs or only effector cells with 50 nM dasatinib were used as negative controls. For the CAR T-cell experiment, the cells were incubated at 37 °C with 5% CO_2_ for 72 hours. One hundred microliters of cell suspension from each well was stained with anti-human CD45 according to the manufacturer’s instructions and washed twice with PBS (Gibco, USA) (300×g, 5 min, 4 °C). The results were evaluated by flow cytometry using ACEA NovoCyte 2060 (Agilent, USA). ACEA NovoExpress (Agilent, USA) was used for data analysis.

#### BM cells

A total of 3×10^5^ BM cells (target cells) were mixed with CD45^Δ^ CAR45 T cells or CD45^Δ^ T cells (E:T ratio 1:3) on the 8th-16th days after CD45 knockout. Wells containing only BM cells or supplemented with 50 nM dasatinib were used as negative controls. For the CAR T-cell experiment, the cells were incubated at 37 °C with 5% CO_2_ for 24, 48 and 72 hours. One hundred microliters of cell suspension from each well was stained with anti-human CD45, CD33, CD3, CD56, CD16, CD19 and 7-AAD according to the manufacturer’s instructions and washed twice with PBS (Gibco, USA) (300×g, 5 min, 4 °C). The results were evaluated by flow cytometry using ACEA NovoCyte 2060 (Agilent, USA). ACEA NovoExpress (Agilent, USA) was used for data analysis.

### Intracellular detection of proinflammatory cytokines

A total of 3×10^5^ CD45^Δ^ T cells (from 2 healthy donors) and T cells (from 2 healthy donors) (effector cells) were mixed with Dynabeads Human T Activator CD3/CD28 (Thermo Fisher Scientific, USA) on the 8th-16th days after knockout of CD45 (E:beads ratio was 1:1) in RPMI 1640 (Gibco, USA) medium without IL-2 and incubated at 37 °C with 5% CO_2_ for 18 hours. Positive control wells containing only effector cells were activated with 40 nM PMA (Invitrogen, USA) and 1 μM ionomycin (Invitrogen, USA) for 3 hours. Wells with untreated effector cells only served as negative controls. After incubation, experimental and control wells were mixed with 10 μg/mL brefeldin A (Sigma, USA) and incubated at 37 °C with 5% CO_2_ for 3 hours. The beads were removed with a DynaMag (Thermo Fisher Scientific), and the cells were washed with PBS (500×g, 5 min, RT). The cells were blocked with 5% FBS (Gibco, USA), stained with anti-human CD45, anti-human CD4 and anti-human CD8a for 15 min at RT, washed with PBS (500×g, 5 min, RT), and fixed with 2% paraformaldehyde (Thermo Fisher Scientific, USA) at 4 °C for 20 min. After fixation, the cells were washed with 0.2% saponin (Merck, Germany) in PBS (500×g, 5 min, RT), blocked with 5% FBS (Gibco, USA), stained with anti-human IFN-γ and anti-human IL2 antibodies for 40 min at RT in 0.2% saponin (Merck, Germany) in PBS (Gibco, USA) and washed twice with PBS (500×g, 5 min, RT). The results were evaluated by flow cytometry using ACEA NovoCyte 2060 (Agilent, USA). Data were analyzed by ACEA NovoExpress software (Agilent, USA). ACEA NovoExpress (Agilent, USA) was used for data analysis. The antibodies used are described in **Supplementary Table 1**.

### In vivo study

Six- to eight-week-old female and male NCG mice with an average weight of 16 to 20 g were used for the experiments.

#### In vivo CAR19 and CD45^Δ^ CAR19 T-cell activity study

Tumors were engrafted by inoculating 1×10^6^ Nalm-6 cells expressing firefly luciferase and GFP intravenously in 100 μL of PBS (Gibco, USA). Mice were randomly assigned to 2 experimental groups and 1 control group. Tumor-bearing mice were subjected to i.v. injection of 5×10^6^ CD45^Δ^ CAR19 T cells, CAR19 T cells (experimental groups), or mock T cells (control group) on the 7th day after tumor engraftment. Tumors were observed weekly for 5 weeks using an IVIS Spectrum In Vivo Imaging System (PerkinElmer, USA) after i.p. injection of D-luciferin (GoldBio, USA) according to the manufacturer’s instructions.

#### In vivo CAR45 and CD45^Δ^ CAR45 T-cell activity study

Tumors were engrafted via *i.v.* inoculation of 2×10^6^ THP-1 cells expressing firefly luciferase and GFP in 100 μL of PBS (Gibco, USA). Mice were randomly assigned to 4 experimental groups and 1 control group. Tumor-bearing mice were subjected to *i.v.* injection of 5×10^6^ CD45^Δ^ CAR45 T cells, CAR45 T cells (experimental groups), or mock T cells (control group) on the 3rd or 7th day after tumor inoculation. Mice that received therapy on the 7th day were injected twice with 2.5×10^6^ CD45^Δ^ CAR45 T cells and CAR45 T cells on the 13th and 20th days. Tumors were observed weekly for 11 weeks, starting on the 6th day after THP-1 injection, using an IVIS Spectrum *In Vivo* Imaging System (PerkinElmer, USA) after i.p. injection of D-luciferin (GoldBio, USA) according to the manufacturer’s instructions.

#### In vivo CAR45 and CD45^Δ^ CAR45 T-cell activity study in hu-PBMC-NCG mice

Mice were subjected to *i.v.* injection of 20×10^6^ PBMCs from Donor 1. After 2 weeks, mice were randomly assigned to experimental and control groups and were injected with 10×10^6^ CD45^Δ^ CAR45 T cells or CD45^Δ^ CAR19 T cells derived from Donor 1. For flow cytometry analysis, blood samples from the retro-orbital sinus were collected. Human CD45+ cell and CAR T-cell counts were monitored by staining of blood samples with anti-human antibodies against CD45, CD3 and IgG (**Supplementary Table 1**), and the results were acquired with a CytoFLEX flow cytometer (Beckman Coulter, USA) in the 2nd, 4th and 8th weeks after PBMC injection.

### Statistics

Statistical analysis was performed using GraphPad Prism 8.0 (GraphPad, USA). Each figure legend denotes the statistical test used. Mean values are plotted as bar graphs, and error bars indicate the standard deviation (SD) unless otherwise stated. ANOVA multiple-comparison p values were calculated using Tukey’s multiple-comparisons test. T test multiple-comparison p values were calculated using the two-stage step-up (Benjamini, Krieger, and Yekutieli) method. All t tests were 2-sided unless otherwise indicated.

### Ethics Approval

All studies were conducted in accordance with protocols approved by the local ethics committee of the Dmitry Rogachev National Medical Research Center of Pediatric Hematology, Oncology, and Immunology (decision on 2018.01.18) and the IBCh RAS Institutional Animal Care and Use Committee. The animals were housed under specific pathogen-free conditions in the Animal Breeding Facility BIBCh RAS (the Unique Research Unit Bio-Model of the IBCh, RAS; the Bioresource Collection – Collection of SPF-Laboratory Rodents for Fundamental, Biomedical, and Pharmacological Studies, Russia), which is accredited at the international level by AAALACi. All procedures were approved by the IBCh RAS Institutional Animal Care and Use Committee.

### Data Sharing Statement

The main data supporting the results in this study are available within the paper and its Supplementary Information. All data generated in this study are available from the corresponding authors on reasonable request.

## Results

### 1. *PTPRC* gene editing of T cells for panhematologic cancer immunotherapy

The expression patterns of CD45 in the bone marrow of patients with various types of acute leukemia make it a promising target in the context of hematological malignancies. CD45 targeting could serve dual purposes, serving as both a conditioning regimen prior to HSCT and a panhematologic cancer immunotherapy (**Figure 1a**). We used CRISPR/Cas9-mediated genome editing to disrupt the *PTPRC* gene (encoding the CD45 protein) in T cells to overcome fratricide-related issues during CD45-specific CAR T (CAR45) cell manufacturing and expansion (**Figure 2a**). We designed different guide RNAs (gRNAs) targeting *exon 1* of the *PTPRC* gene to prevent translation of all possible isoforms of CD45 and tested them in activated human T cells. For further experiments, we chose a gRNA variant (gRNA-2; **Supplementary Figure 1a**) that disrupted the *PTPRC* gene with high efficiency, as evidenced by the loss of surface CD45 expression in >85% of T cells four days post electroporation (**Figure 1b and c; Supplementary Figure 1b**). The knockout of CD45 did not result in significant changes in T cell viability (**Figure 1d)**, with stable CD45 ablation for more than 4 weeks of cell culture **(Supplementary Figure 1c)**. Loss of CD45 expression was further confirmed by confocal microscopy and Western blot analysis of cells with disrupted *PTPRC* gene expression after FACS (**Figure 1e and f, Supplementary Figure 1d**). Analysis of the targeted fragment in PCR-amplified genomic DNA by Sanger sequencing demonstrated the presence of indels in the CD45 gene in CD45^Δ^ T cells **(Supplementary Figure 1e)**. Then, to explore the safety of the designed *PTPRC* gene editing approach, potential off-target sites for gRNA-2 were verified by next-generation sequencing of gene-edited T-cell genomic DNA. The percentage of reads containing insertions and deletions increased from 70 to 90 with the increase in knockout percentage over time exclusively in CD45^Δ^ T cells (4, 8 and 16 hours following electroporation with Cas9/gRNA-2) **(Supplementary Figure 1f)**. For gRNA-2, only one off-target site with maximal predicted disruption probability has higher percent of both insertions and deletions for the cells treated with Cas9/guide RNA complex compared to a negative control at specified timepoints after the electroporation. It was determined as the *THSD4* gene intron (0.5%; P<.001), but the *THSD4* gene was not expressed in immune cells **(Supplementary Table 2)**. These results suggest that Cas9/gRNA-2-mediated disruption of the *PTPRC* gene 1 exon generates viable CD45^Δ^ T cells with ablated cell surface expression of CD45.

**Figure 1.**
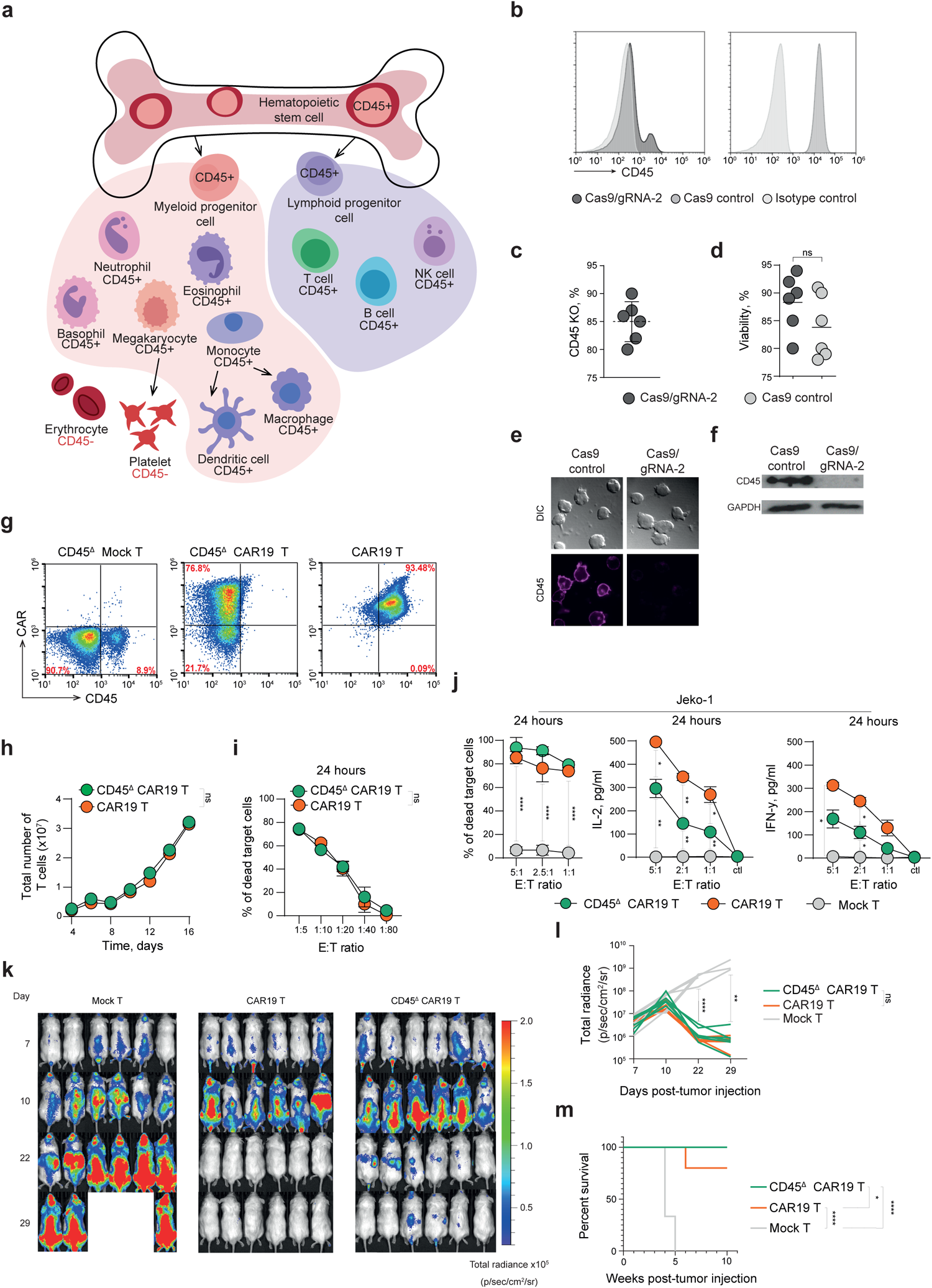
Disruption of CD45 expression does not impair the functional activity of T cells and CAR T cells. **a.** CD45 expression on different cells during hematopoiesis. **b.** Representative histogram showing ablation of CD45 expression in T cells after electroporation with CRISPR/cas9 and CD45-specific gRNA-2. **c, d**. Frequency and viability of CD45-negative cells on the 4th day after electroporation with Cas9/gRNA-2 complexes. P values were determined by multiple unpaired t tests. Each dot represents an independent donor. **e.** Fluorescence microscopy of T and CD45^Δ^ T cells stained with anti-CD45 antibodies (magenta) after FACS. **f.** Western blot analysis of CD45 expression in T cells from **b** after FACS. **g.** Representative dot plots showing the expression of CAR19 and CD45 in mock-transduced, CAR19 and CD45 ^Δ^ CAR19 T cells. **h.** Total expansion of CAR19 and CD45^Δ^ CAR19 T cells after 14 days of *in vitro* culture. P values were determined by multiple unpaired t tests. **i.** Nalm-6 cells were incubated with different numbers of CAR19 and CD45^Δ^ CAR19 T cells prior to analysis of tumor cell lysis. P values were determined by multiple unpaired t tests. **j.** IL-2 and IFN-γ secretion and target cells killing by mock-transduced, CAR19 and CD45^Δ^ CAR19 T cells mixed with Jeko-1 cells at different effector (E)-to-target (T) ratios. P values were determined by multiple unpaired t tests. Nonsignificant values are not shown. **k.** Representative IVIS images of mice from the mock-transduced, CAR19 and CD45^Δ^ CAR19 T-cell-treated groups. NCG mice were implanted i.v. with 1×10^6^ Nalm-6/ffluc cells. On day 7 after tumor inoculation, the animals were subjected to *i.v.* infusion of 3×10^6^ mock-transduced, CAR19 or CD45^Δ^ CAR19 T cells (n=6). **l.** Quantification of tumor burden (as the total radiance due to luciferase activity per mouse) from **k** for days 7-29. P values were determined by multiple unpaired t tests. **m.** Survival plots for animals from the experimental and control groups. Overall survival curves were plotted using the Kaplan‒Meier method and compared using the log-rank (Mantel‒Cox) test. Data from **b, e-j** represent independent replicated experiments with cells isolated from 3 donors. All data represent the mean ± SD.

**Figure 2.**
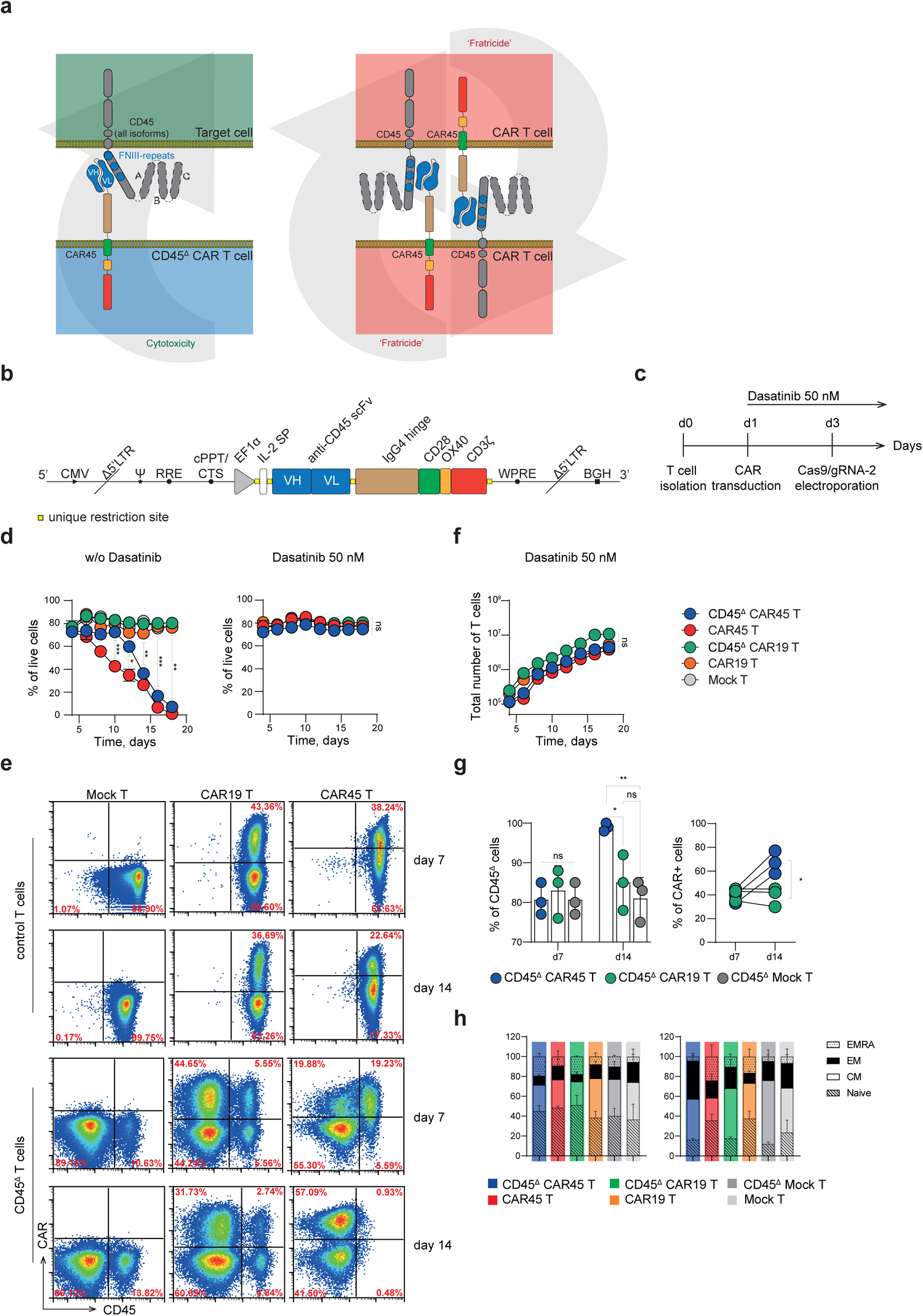
Generation of fratricide-resistant CD45-specific CAR T cells. **a.** Experimental setup for CD45^Δ^ CAR T-cell engineering. T cells were electroporated with Cas9/gRNA-2 complexes and transduced with CAR. **b.** Schematic of CAR45 construct design. **c.** Schematic timeline of CD45^Δ^ CAR T-cell engineering. **d.** Effect of 50 nM dasatinib on the viability of CAR19, CAR45, CD45^Δ^ CAR19, and CD45^Δ^ CAR45 T cells in culture. P values were determined by multiple unpaired t tests. **e.** Representative dot plots showing the expression of CD45 and CARs in control and Cas9/gRNA-2-electroporated T cells on days 7 and 14 after transduction. **f.** Total expansion of cells from **d, right**. P values were determined by multiple unpaired t tests. **g.** Percentage of CD45^Δ^ CAR T cells in **e**. P values were determined by multiple unpaired t tests. **h.** Differentiation state analysis of CAR19-, CAR45- and mock-transduced T cells electroporated with Cas9/gRNA-2. Cells were stained with anti-CD44 and anti-CD62L antibodies and analyzed by flow cytometry on days 7 and 14 after transduction. CM - central memory, EM - effector memory, EMRA - terminally differentiated T cells. P values were determined by two-way ANOVA. Data from **d-h** represent independent experiments with cells isolated from 5 donors. All data represent the mean ± SD.

### 2. Genetic disruption of the CD45 gene does not affect T- or CAR T-cell functionality

Then, we questioned whether *PTPRC* gene disruption and the absence of CD45-related phosphatase activity influence T-cell functionality. First, we compared the cytotoxicity and cytokine release of T and CD45^Δ^ T cells in response to the Ramos cell line with a bispecific CD19/CD3 T-cell engager (BiTE) **(Supplementary Figure 2a)**. Interestingly, T and CD45^Δ^ T cells displayed equal cytotoxic activity, although less IL-2 and IFN-γ was secreted by CD45^Δ^ T cells **(Supplementary Figure 2b, 2c)**. These results suggest that CD45^Δ^ T cells remain cytotoxic and secrete inflammatory cytokines.

The high cytotoxic efficiency of CD45^Δ^ T cells encouraged us to evaluate the impact of *PTPRC* gene disruption on CAR T-cell performance. First, we electroporated T cells with CRISPR/Cas9 and CD45-specific gRNA-2 or control gRNA and transduced them with CD19-specific CAR19 (**Figure 1g)** because the activity of CAR19 is well documented both *in vitro* and *in vivo*. In comparison to regular CAR19 T cells stimulated under the same conditions, loss of CD45 did not reduce the proliferation rate of CD45^Δ^ CAR19 T cells (**Figure 1h).** Then, we compared the cytotoxicity of regular CAR19 T cells and CAR19 T cells electroporated with Cas9/gRNA-2. CAR19 and CD45^Δ^ CAR19 T cells were cocultured with Nalm-6 and Jeko-1 cells expressing CD19. Both CAR19 and CD45^Δ^ CAR19 T cells efficiently eliminated tumor cells following 24 hours of coculture (**Figure 1i, j left)**. In line with that finding, compared to CAR19 T cells, CD45^Δ^ CAR19 T cells secreted lesser but comparable levels of IL-2 and IFN-γ in response to incubation with target cells (**Figure 1j middle and right)**.

To explore the cytotoxic function of CD45^Δ^ CAR19 T cells *in vivo*, we employed a mouse xenograft model of pre-B-cell lymphoma with Nalm-6 cells. NOD–severe combined immunodeficiency (SCID) γ chain-deficient (NCG) mice were intravenously engrafted with Nalm-6/ffluc cells expressing firefly luciferase. On day 7 following Nalm-6 cell injection, when the tumors could first be detected, each recipient mouse was intravenously injected with a single dose of 2х10^6^ CD45^Δ^ CAR19, CAR19 or mock-transduced T cells. *In vivo* imaging of animals revealed a significant reduction in tumor burden in the CAR19 and CD45^Δ^ CAR19 therapy groups (**Figure 1k)**. Additionally, we found no differences in tumor growth kinetics between the groups of mice treated with regular or CD45^Δ^ CAR19 T cells (**Figure 1l)**. The reduction in tumor burden led to high survival rates after injection of CD45^Δ^ CAR19 T cells and slightly worse rate in group of mice treated with regular CAR19 T cells (**Figure 1m)**. Therefore, CD45 knockout had no negative effect on the short-term persistence or cytotoxicity of CAR T cells in our mouse tumor xenograft model.

### 3. Generation of fratricide-resistant CD45-specific CAR T cells

Shared CD45 expression between CAR T cells and target cells represents a significant challenge for the development of CAR T-based strategies to target CD45+ cells. After obtaining experimental proof that CD45^Δ^ CAR19 T cells possess potent antitumor activity and persistence in mice, we next explored whether CD45 knockout CAR T cells targeting CD45 have favorable biological characteristics (**Figure 2a)**. We generated a CAR45 construct by fusing a CD45-specific single-chain variable fragment derived from the antibody clone BC8 to a third-generation CAR backbone containing an IgG4 CH2-CH3 spacer and cytoplasmic endodomains from CD28, CD134 and CD3ζ^42,45^ (**Figure 2b)**. Knockout of CD45 was performed 2 days following transduction of T cells with the CAR constructs (**Figure 2c)**. However, in striking contrast to the control CD45^Δ^ CAR19 T cells, both CD45^Δ^ CAR45 and CAR45 T cells exhibited poor viability and failed to expand (**Figure 2d, left)**. Presumably, the expression of trace amounts of CD45 on T cells led to antigen-induced fratricide and chronic stimulation mediated by CAR45. The expression of CAR45 and CD45 was verified on days 7 and 14 post CD45 knockout (**Figure 2e)**. Inspired by previous studies showing that fratricide of CAR T cells can be fully prevented using dasatinib, the pharmacologic inhibitor of key CAR/CD3ζ signaling kinases^46^, we supplemented the T cell culture with 50 nM dasatinib, which rescued the expansion and viability of CAR45 and CD45^Δ^ CAR45 T cells (**Figure 2d right and f).** Interestingly, in the culture of CD45^Δ^ CAR45 T cells, despite the inclusion of dasatinib, we observed the elimination of residual CD45-positive cells and the growth of the CD45^Δ^ CAR45 population on day 14 (**Figure 2g).** Removal of CD45 did not alter the phenotype of CD45^Δ^ CAR19 or CD45^Δ^ CAR45 T cells. On day 14 of cultivation, central memory (Tcm) and effector memory (Tem) cells predominated among CD45^Δ^ CAR19, CD45^Δ^ CAR45 and mock-transduced CD45^Δ^ T cells (**Figure 2h)**.

### 4. CD45^Δ^ CAR45 T cells outperform CAR45 T cells *in vitro* and *in vivo*

To assess whether CD45^Δ^ CAR45 T cells possess functional fratricide-resistance, we analyzed the specificity and cytotoxicity of CD45^Δ^ CAR45 T cells against CD45-positive T-ALL (Jurkat), B-ALL (Jeko-1), AML (THP-1) cells and a CD45-negative (Nalm-6) tumor cell line (**Figure 3a)**. We used primary activated T cells isolated from the same donors to generate CAR19, CAR45, CD45^Δ^ CAR19 and CD45^Δ^ CAR45 T cells. Then, we cocultured CAR T cells with a panel of hematological tumor cell lines with varying expression of CD19 and CD45. As expected, CAR19 and CD45^Δ^ CAR19 T cells specifically eliminated CD19-positive Nalm-6 and Jeko-1 cells with equal efficacy (**Figure 3b)** and similar dynamics (**Figure 3c)**. However, we observed a significant difference in cytotoxicity and cytokine release between CAR45 and CD45^Δ^ CAR45 T cells. In comparison with CAR45 T cells, CD45^Δ^ CAR45 T cells were more effective against CD45^+^ Jurkat, Jeko-1, and THP-1 cells and were not cytotoxic to control CD45-negative Nalm-6 cells, indicating CD45-specific cell killing (**Figure 3b)**. Moreover, when we compared the dynamics of cell killing at a low effector:target ratio (1:3), CD45^Δ^ CAR45 T cells eliminated tumor cells faster than CAR19 and CD45^Δ^ CAR19 T cells (**Figure 3c)**. We also observed that CAR45 T cells exhibited nonspecific hyperactivation, accompanied by elevated production of IL-2 and IFN-γ, even in the absence of target cells and independent of the effector:target ratio. In addition, CD45^Δ^ CAR45 T cells released cytokines only upon incubation with target cells in a manner dependent on the effector:target ratio (**Figure 3d)**.

**Figure 3.**
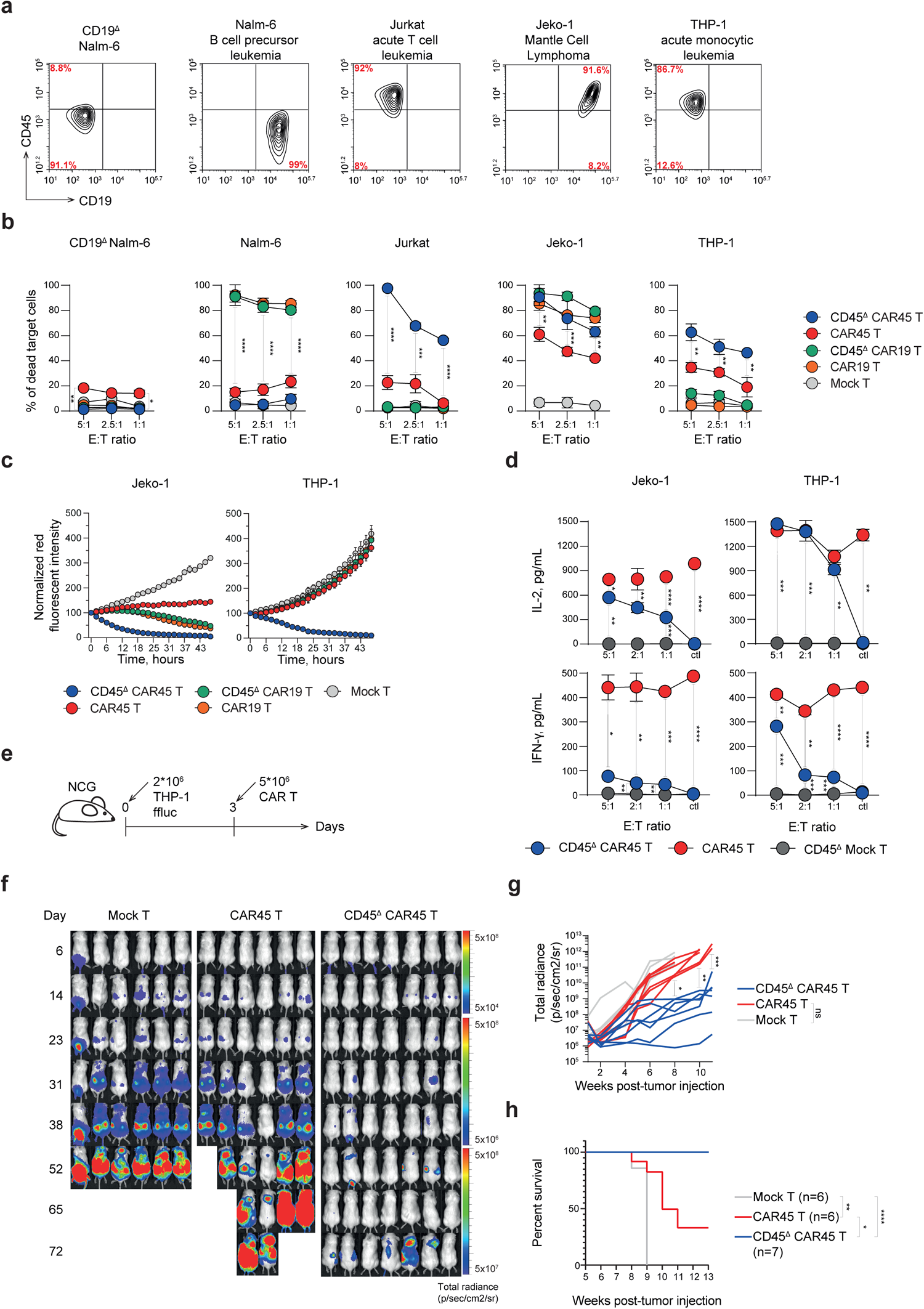
CD45^Δ^ CAR45 T cells outperform CAR45 T cells *in vitro* and *in vivo*. **a.** Representative density plots showing the expression of CD19 and CD45 in leukemia and lymphoma cell lines. **b.** CAR19, CAR45, CD45^Δ^ CAR19, and CD45^Δ^ CAR45 T cells were incubated with blood cancer cell lines prior to tumor cell lysis analysis. Total numbers of live tumor cells were quantified by flow cytometry at 24 hours using counting beads. P values were determined by multiple unpaired t tests. Nonsignificant values are not shown. **c.** Real-time detection of fluorescent Jeko-1 and THP-1 targets incubated with the designed CAR19, CAR45, CD45^Δ^ CAR19, and CD45^Δ^ CAR45 T cells in the IncuCyte killing assay (E:T ratio 1:3). **d.** Quantification of IFN-γ and IL-2 secretion by CAR45 and CD45^Δ^ CAR45 T cells cocultured with Jeko-1 and THP-1 cells. P values were determined by multiple unpaired t tests. **e.** NCG mice were subjected to *i.v.* infusion of 2×10^6^ THP-1/ffluc cells. On day 3 after tumor inoculation, animals were subjected to *i.v.* infusion of 3×10^6^ mock-transduced, CAR45 or CD45^Δ^ CAR45 T cells. **f.** Representative IVIS images of mice from the mock-transduced, CAR45 and CD45^Δ^ CAR45 T-cell-treated groups. **g.** Quantification of tumor burden (as the total radiance due to luciferase activity per mouse) from **f.** for days 7-29. P values were determined by multiple unpaired t tests. **h.** Kaplan–Meier curve showing overall animal survival in each experimental group. P values were determined by the log-rank Mantel‒Cox test. Data from **b-d** represent independent experiments with cells isolated from 3 donors. All data represent the mean ± SD.

Following these *in vitro* tests of T-cell activity, we analyzed the therapeutic efficacy of CD45^Δ^ CAR45 T cells in a mouse xenograft model of disseminated monocytic AML. NCG mice were engrafted via intravenous injection of 2×10^6^ THP-1/ffluc cells expressing firefly luciferase (**Figure 3e)**. Three days after tumor engraftment, the animals were intravenously injected with 5×10^6^ Mock, CAR45 or CD45^Δ^ CAR45 T cells. *In vivo* visualization of animals from the control and therapeutic groups revealed the outstanding therapeutic potential of the CD45^Δ^ CAR45 T cells against disseminated monocytic AML (**Figure 3f)**. All animals treated with CD45^Δ^ CAR45 T cells exhibited tumor elimination and high survival rates (**Figure 3g and h)** (63 days in mock T-cell-treated vs. >90 days in the CD45^Δ^ CAR45 T group; P < 0.001 by the Mantel‒Cox log-rank test). In a parallel experiment, animals subjected to *i.v.* injections of 2.5×10^6^ CD45^Δ^ CAR45 T cells on days 7, 13 and 20 exhibited tumor elimination and extended survival **(Supplementary Figure 3a-d)**, although this regimen was not as effective as the single-dose CAR T-cell injection at a higher dose.

### 5. CD45^Δ^ CAR45 T and NK cells deplete human hematopoietic cells *in vitro*

To evaluate the use of CD45^Δ^ CAR45 T cells to deplete not only malignant cells but also endogenous hematopoietic cells, we incubated primary human T cells and PBMCs with CD45^Δ^ CAR45 T cells. After 24 hours of coculture, we observed that CD45^Δ^ CAR45 T cells but not control CD45^Δ^ CAR19 T cells exhibited killing of T cells and PBMCs (**Figure 4a)**. To assess the kinetics of hematopoietic cell depletion, we mixed PBMCs and CD45^Δ^ CAR45 T cells at a 1:3 ratio and analyzed changes in the cell population for the next 24, 48 and 72 hours (**Figure 4b)**. A negative control group was supplemented with 50 nM dasatinib. We observed complete elimination of PBMCs in the experimental group after 72 hours of coculture, while the PBMCs in the control group remained viable throughout the cultivation period (**Supplementary Figure 4**). To estimate the activity of CAR45 against bone marrow cells, we mixed CD45^Δ^ CAR45 T cells and freshly isolated autologous donor BM cells at a 1:3 ratio and analyzed the changes in cell subsets for the next 24, 48 and 72 hours and cytotoxicity **(Supplementary Figure 5)**.

**Figure 4.**
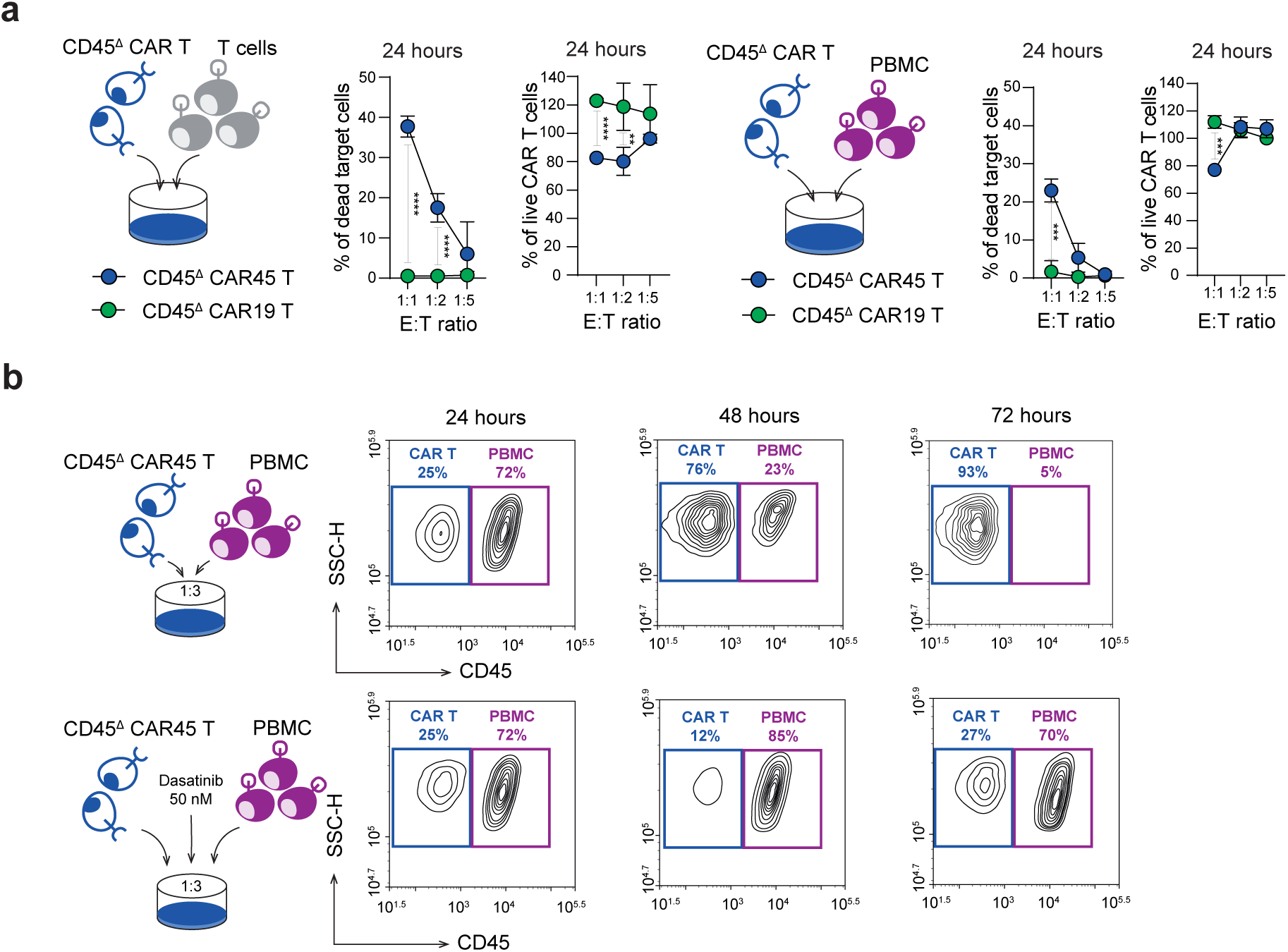
Cytotoxicity and specificity of CD45^ι1^ CAR45 T cells against primary human cells. **a.** Freshly isolated human T cells or PBMCs were cocultured with CD45^ι1^ CAR19 and CD45^ι1^ CAR45 T cells. Total numbers of live target cells were quantified by flow cytometry at 24 hours using counting beads. P values were determined by multiple unpaired t tests. Nonsignificant values are not shown. **b.** Freshly isolated PBMCs were cocultured with CD45^ι1^ CAR45 T cells at a 1:3 ratio for 24, 48 and 72 hours upon addition of 50 nM dasatinib or no drug. Contour plots show the frequency of live PBMCs at the end of the coculture period. Data represent independent experiments with cells isolated from 3 donors. CAR NK cells are considered strong alternatives to traditional CAR T cells due to their potent cytotoxicity and low risk of alloreactivity^47^. Another advantage of CAR NK cells is their short lifespan compared to CAR T cells, which makes them suitable for transient short-term immunotherapy^48^. Therefore, the feasibility of generating functional fratricide-resistant CD45^Δ^ CAR45 NK cells was tested in our study. The protocol used for *PTPRC* gene disruption and CAR transduction was optimized for human NK cells (Figure 5a). CD45 knockout was highly effective and did not alter NK viability or proliferation, and the CAR45 transduction efficacy was higher than 50% (Figure 5b). CD45^Δ^ CAR45 NK cells were expanded in cultures supplemented with 50 nM dasatinib to avoid undesired fratricidal effects. Similar to T cells, CD45^Δ^ CAR45 NK cells demonstrated outstanding killing of allogeneic and autologous T cells and PBMCs following 4 and 24 hours of incubation (Figure 5c). These results support the potential implementation of fratricide-resistant CD45^Δ^ CAR45 NK cells for adoptive immunotherapy.

**Figure 5.**
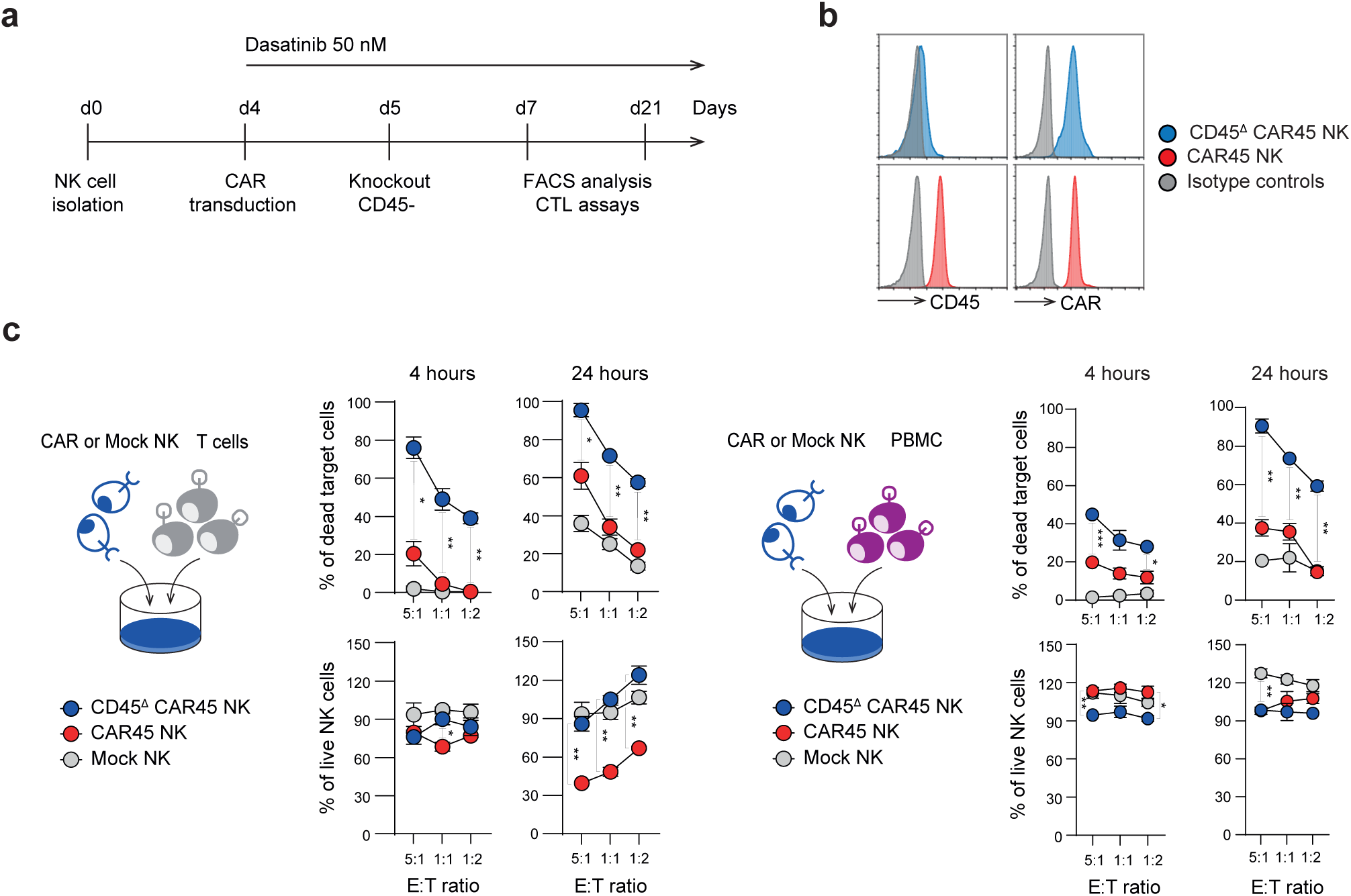
Cytotoxicity and specificity of CD45^Δ^ CAR45 NK cells against primary human cells. **a.** Schematic of the CD45^Δ^ CAR NK cell engineering timeline. **b.** Representative dot plots showing the expression of CD45 and CAR45 in control and Cas9/gRNA-2-electroporated NK cells on day 14 after transduction. **c.** Freshly isolated autologous human T cells or PBMCs were cocultured with CAR45 and CD45^Δ^ CAR45 NK cells. Total numbers of live CAR and target cells were quantified by flow cytometry at 4 and 24 hours. P values were determined by multiple unpaired t tests. Nonsignificant values are not shown. Data from **b-c** represent independent experiments with cells isolated from 3 donors. All data represent the mean ± SD.

### 6. Autologous CD45^Δ^ CAR45 T cells target and deplete hematopoietic cells in humanized hu-PBMC-NCG mice

We next determined whether conditioning with CD45^Δ^ CAR45 T cells enabled successful depletion of engrafted donor hematopoietic cells in humanized hu-PBMC-NCG mice. Human PBMC-engrafted NCG mice (hu-PBMC-NCG) were generated by transferring 20×10^6^ PBMCs from healthy donors into NCG mice (**Figure 6a)**. Four weeks after PBMC engraftment, double staining of blood samples with anti-human and anti-mouse CD45 antibodies was performed to estimate chimerism in the animals (**Figure 6b)**. Successful engraftment occurred in all animals, with 40-60% of the cells of human origin four weeks after PBMC injection. The humanized mice were split randomly into three groups (n=8) and subjected to *i.v.* injection of 10×10^6^ CD45^Δ^ CAR45, CD45^Δ^ CAR19, or CD45^Δ^ mock-transduced T cells. Mice treated with CD45^Δ^ CAR45 T cells exhibited decreasing counts of engrafted human PBMCs four weeks after injection. The disappearance of human PBMCs was recorded in week eight (**Figure 6b and d)**. In contrast, animals injected with CD45^Δ^ CAR19 or CD45^Δ^ mock-transduced T cells maintained stable or even increasing levels of humanization. Analysis of the persistence of human cells in the blood showed the consistent presence of only CD45^Δ^ CAR45 T cells, which accounted for approximately 25% of total CD3+ T cells (**Figure 6c and e)**. Typically, hu-PBMC-NCG mice develop a xenogeneic response resembling GvHD 4-5 weeks following human PBMC injection. This is consistent with the survival time of animals treated with CD45^Δ^ CAR19 or CD45^Δ^ mock-transduced T cells. However, the effective elimination of hPBMCs by CD45^Δ^ CAR45 T cells dramatically delayed the development of GvHD-like symptoms and increased the survival of the humanized mice by forty days (65 days vs. 105 days, respectively; p=.0092) (**Figure 6f)**.

**Figure 6.**
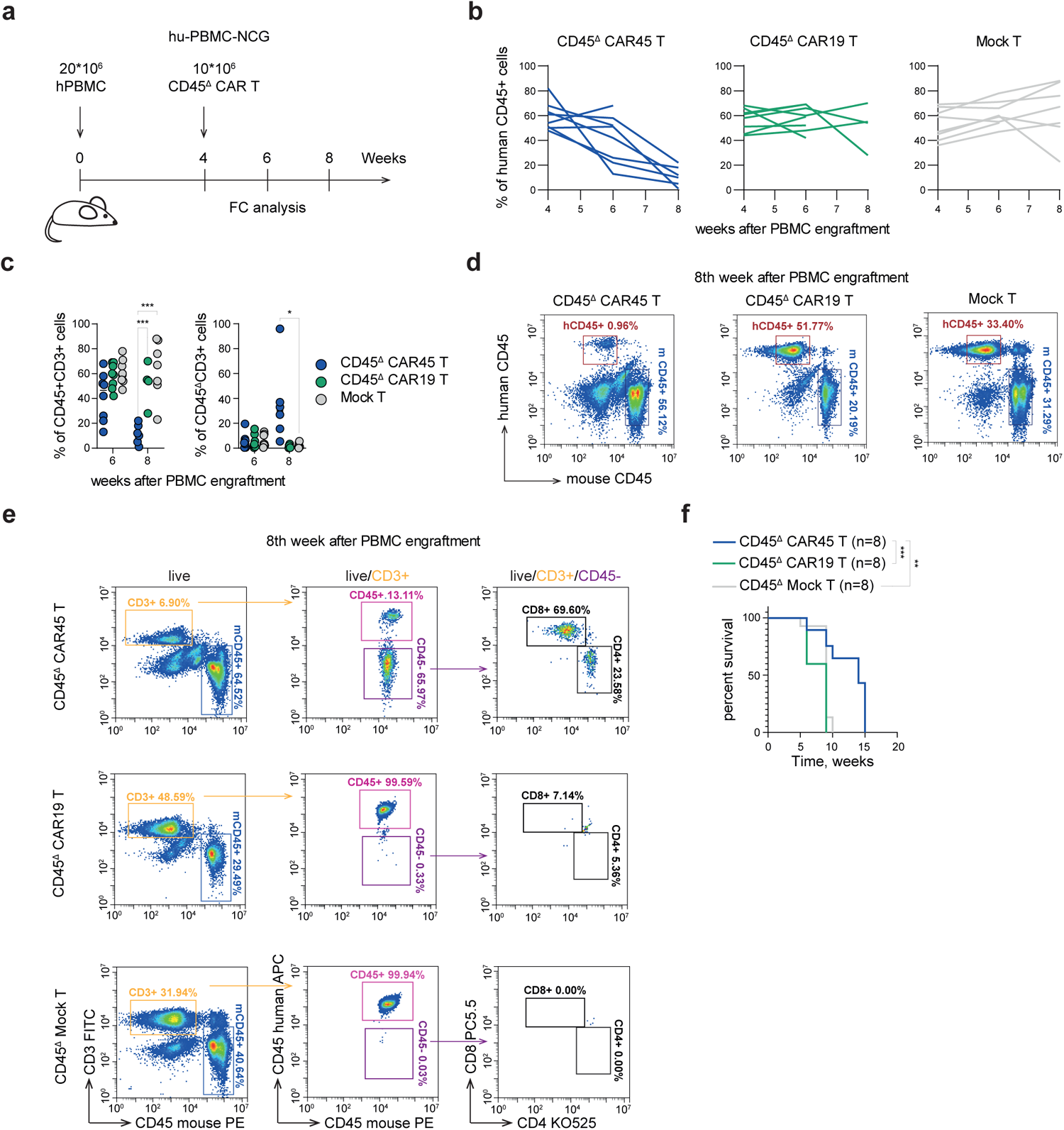
CD45^Δ^ CAR45 T cells eliminate healthy autologous human PBMCs *in vivo*. **a.** Schematic of the mouse model setup. NCG mice were subjected to i.v. infusion of 20×10^6^ human PBMCs. On day 4 after PBMC engraftment, the animals were subjected to *i.v.* infusion of 10×10^6^ CD45^Δ^ CAR19, CD45^Δ^ CAR45 and CD45^Δ^ mock-transduced T cells. **b.** Effect of CD45^Δ^ CAR45 T cells on autologous human PBMCs *in vivo*. Hu-PBMC-NCG mouse chimerism was evaluated by flow cytometry on weeks 4, 6 and 8 after PBMC engraftment (as the percentage of circulating human CD45-positive cells). Each line represents individual mice. **c.** Number and percentage of circulating CD45^-^/CD3^+^ human cells analyzed at weeks 6 and 8 of the experiment. Each dot represents an individual mouse. P values were determined by multiple unpaired t tests. Nonsignificant values are not shown. **d.** Representative dot plots of the chimerism analysis for the mice in **b** on week 8 of the experiment. Numbers indicate the percentages of human CD45 (hCD45) and mouse (mCD45) cells in the sample. **e.** Gating strategy and representative dot plots of flow cytometry analysis of CAR T-cell persistence at week 8 post injection. **f.** Kaplan–Meier curve showing overall animal survival in each experimental group. P values were determined by the log-rank Mantel‒Cox test. Nonsignificant values are not shown. All data represent the mean ± SD.

It is important to acknowledge the limitations of the hu-PBMC-engrafted animal model. The hu-PBMC-NCG model allowed us to evaluate the *in vivo* efficacy of CD45^Δ^CAR45 T cells against healthy primary human cells. However, only additional *in vivo* studies on intact murine immune system-conditioned murine anti-mouse CD45 CAR T cells will allow us to assess potential trafficking issues, which were previously reported to reduce the efficacy of murine CD117-CAR T cells^18^ but not human CD117-CAR T cells in NSG mice engrafted with AML patient BM cells^19^.

## Discussion

A critical consideration in employing the CAR T cell approach or antibody-drug conjugate for BM conditioning lies in the deliberate avoidance of ionizing radiation and potent alkylating agents such as busulfan. Such agents have the potential to cause damage or inflammation to both the BM and thymic stroma, thereby creating an environment that may be less conducive to successful BM engraftment and eventual restoration of lymphoid immunity^49^.

We provide the first proof-of-concept demonstration of the generation and preclinical assessment of fratricide-resistant CAR T-cells directed to target CD45-expressing human hematopoietic cells (referred to as CD45^Δ^CAR45 T cells). We show that the developed CRISPR/Cas9 CD45 gene disruption and CAR transduction protocol consistently yields a highly enriched population of gene-edited CD45^Δ^CAR45 T and NK cells; CD45 gene disruption does not compromise cytotoxicity or functional potency of T cells. CD45^Δ^CAR45 T cells specifically kill CD45+ AML, B-ALL, T-ALL, and MCL cell lines and primary HSPCs; CD45^Δ^CAR45 T cells deplete AML cell lines as well as healthy donor PBMCs *in vivo* in mouse xenograft models. These findings establish CD45 as a potential candidate for CAR-mediated immunotherapy against leukemia, and, simultaneously, as a preconditioning strategy before HSCT.

A panel of antigens, including CD117, CD123, CD33, and others, are considered potential targets for HSCT conditioning. Data on an anti-CD117 antibody and an ADC were recently reported, and early clinical experience suggests that they have high potential as engraftment-facilitating agents^50,51^. Notably, monotherapy with anti-CD117 might not provide sufficient immune suppression for the engraftment of allogeneic hematopoietic cells beyond the cohort of severe combined immune deficiency patients. The same is true for other potential myeloid-directed immunotherapeutics, as they will, by definition, spare the lymphoid compartments. Moreover, it has been suggested that anti-CD117 antibodies and similar myeloid-targeted agents can be combined with broadly immunosuppressive agents, including CD45-targeted agents^52,53^, to achieve the goal of HSCT conditioning.

A particularly attractive target for hematologic malignancies and HSCT for nonmalignant indications is CD45, a common leukocyte antigen expressed by almost all nucleated white cells, including T cells, NK cells, and granulocytes; the level of CD45 expression generally increases as the cells mature. CD45 targeting was pioneered by M. Brenner^16^; unfortunately, the naked antibody did not reach the late clinical trial stage. For over two decades, anti-CD45 radioconjugates were developed, mostly aimed at leukemia and lymphoma patients^54–56^. Ample preclinical and early clinical evidence demonstrated that this approach could deliver high doses of radiation with high efficacy and limited direct toxicity^57–59^. Using radioconjugates has obvious limitations in cohorts of patients with nonmalignant disorders and in children, where long-term genotoxicity should be minimized. The use of an anti-CD45-based ADC was recently proposed as an approach to nongenotoxic conditioning for patients with primary immune deficiency and constitutional bone marrow failure^10,12,60^. The results of this promising preclinical work are awaiting confirmation in human studies.

CD45 targeting by CAR T cells has not been considered feasible for several reasons. (1) shared antigen expression between the target and CAR T cells can result in fratricide. Only recent studies that developed gene-editing approaches to generate CD5- and CD7-specific CAR-T cells demonstrated the possibility of obviating fratricide. Of note, naturally CD5-negative and CD7-negative T cells can be selected to produce CAR T cells of the relevant specificity, a phenomenon not described for CD45^31,36,46^. (2) Considering the severe combined immune deficiency phenotype generated by human constitutional CD45 deficiency^61^, the idea that the removal of such an important molecule as CD45 will retain T-cell functionality seems counterintuitive. In addition, the importance of CD45 expression for T cells, driven by synthetic CARs, was not clearly established.

Recently an elegant alternative approach to target CD45 was reported, based on base-editing of HSC to eliminate the epitope targeted by BC8 CAR^62^.

There are multiple ways to further improve CD45^Δ^CAR45 for clinical implementation. This approach could be tested in the autologous setting or further developed with multiple gene edits^63^ into an allogeneic or even universal, “off-the-shelf” product. Since long-term persistence is not the desired characteristic of the CD45 targeting agent, safety switches, markers for antibody-mediated depletion, transient CAR expression via mRNA or, alternatively, adapter molecule-based CAR design could be considered.

In summary, we established gene-edited anti-human CD45 CAR T cells as a highly effective immunotherapeutic modality for leukemia, and we envision that this approach will increase the curative potential of preconditioning regimens prior to allogeneic HSCT. For nonmalignant indications, it would provide the necessary immune suppression and create “space” for engraftment. In the case of hematologic malignancies, it would exert additional anti-leukemia activity. Alternatively, CAR45 could be used as an add-on anti-leukemia agent with a nonoverlapping toxicity profile and resistance mechanisms in cases of refractory leukemia.

## Authorship Contributions

V.M.S., D.V.V.,D.S.O., W.W., Y.H., D.E.P., M.S.F., E.A.M., E.A.K., D.Z., I.Z.M., A.S.C., Y.P.R., A.V.S. performed experiments, analyzed, and interpreted data; L.C., Z.M., H.Z., J.X., A.G.G., P.W., I.Z.M., G.B.T., Y.P.R., A.G.G., M.A.M, and A.V.S. analyzed and interpreted data and revised the manuscript; P.W., M.A.M, and A.V.S. designed research, analyzed, and interpreted data, and wrote the paper.

