## Supplemental Figures, Tables for "Targeting CD45 by gene-edited CAR-T cells for leukemia eradication and hematopoietic stem cell transplantation preconditioning"

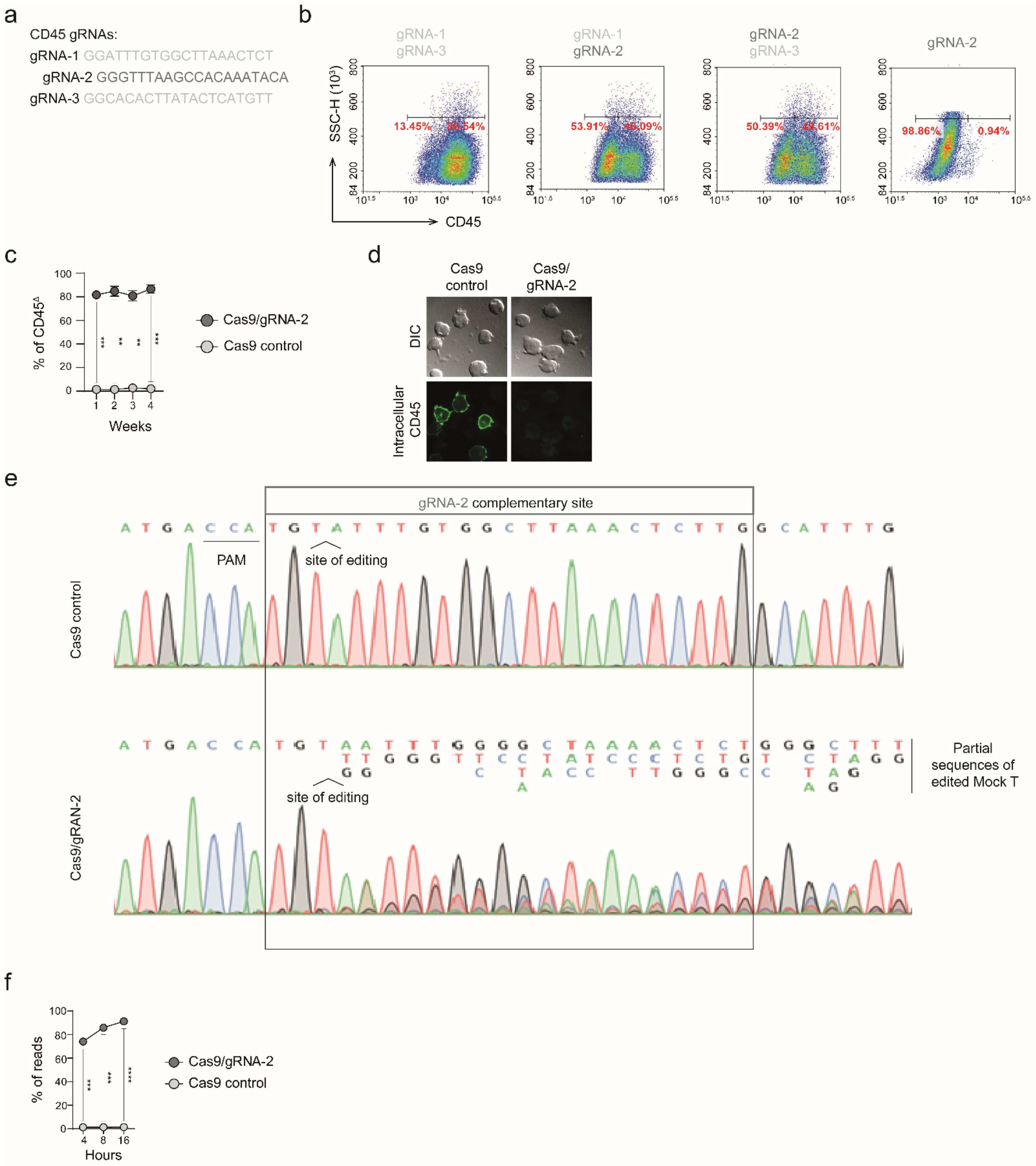


**Supplementary figure 1. Design and validation of gRNAs for CRISPR/Cas9-mediated *PTPRC* gene disruption.**

**a.** gRNA target sequences located in the exon 1 sequence of the *CD45 gene.* Dark gray indicates the selected gRNA.

**b.** T cells were electroporated with different combinations of 2 out of 3 gRNAs, and knockout efficiency was analyzed by flow cytometry. Representative dot plots are shown.

**c.** Analysis of the stability of the CD45$^{\Delta}$ T-cell population during 4 weeks of cultivation. P values were determined by multiple unpaired t tests.
**d.** Intracellular (green) detection of CD45 expression on T and CD45$^{\Delta}$ T cells. Representative images are shown.
**e.** Sanger sequencing of the *CD45 gene* in CD45$^{\Delta}$ T cells.
**f.** Number of indels in the *CD45 gene* after 4, 8 and 16 hours of electroporation with gRNA2. Cells electroporated with Cas9 protein only were used as a negative control. Data are shown as the mean ±SD and represent two independent experiments. P values were determined by multiple unpaired t tests.
Data from **b-d** represent independent experiments with cells isolated from 3 donors.
All data represent the mean ± SD.

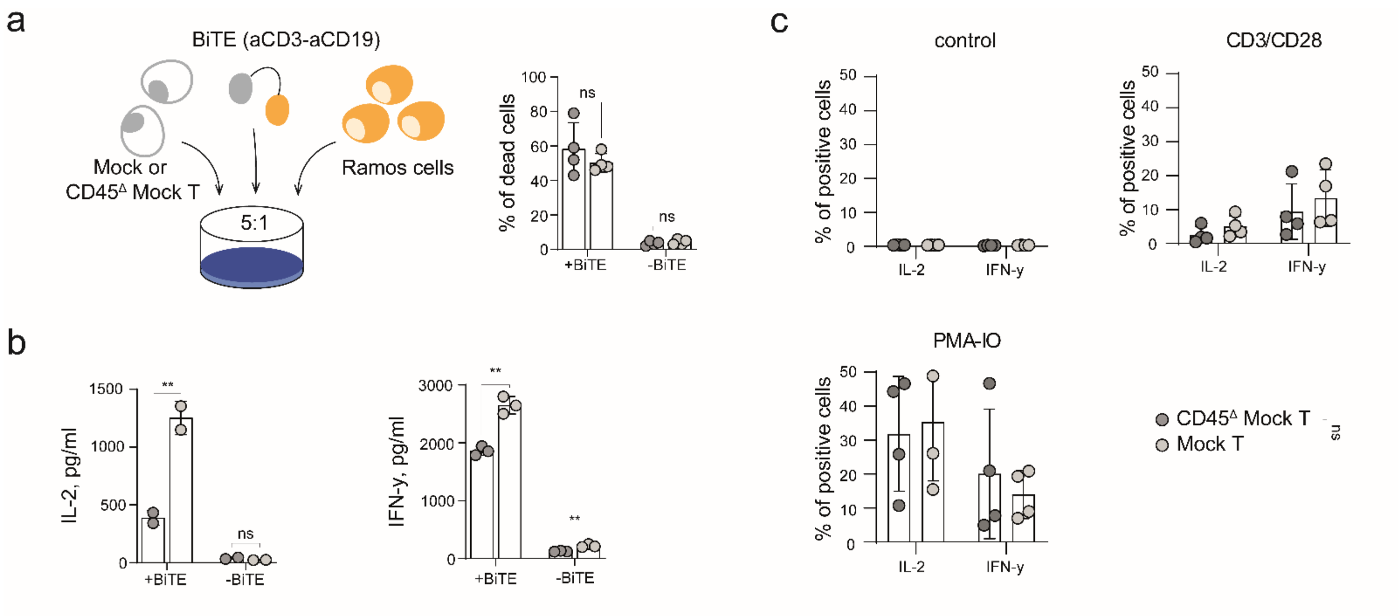


**Supplementary figure 2. Analysis of the functional activity of CD45$^{\Delta}$ T cells after electroporation with Cas9/gRNA-2 complexes.**

**a.** Cytotoxicity of human T cells and CD45$^{\Delta}$ T cells mixed with target CD19$^{+}$ Ramos cells at a 5:1 ratio in the presence of 1 nM BiTE blinatumomab (anti-CD3-anti-CD19). Human CD45$^{\Delta}$ T cells were sorted to 100% prior to the experiment and incubated with Ramos cells for 24 hours. P values were determined by multiple unpaired t tests.

**b.** Comparison of IL-2 and IFN-γ cytokine secretion by normal and CD45$^{\Delta}$ T cells after electroporation with Cas9/gRNA-2 complexes. T cells and CD45$^{\Delta}$ T cells were cocultured with Ramos cells at a 5:1 E:T ratio in the presence of 1 nM BiTE blinatumomab (anti-CD3-anti-CD19). P values were determined by multiple unpaired t tests.

**c.** Flow cytometry analysis of intracellular concentrations of IL-2 and IFN-γ in human T cells stimulated with anti-CD3/CD28 Dynabeads (Invitrogen) or PMA/IO for 18 hours. P values were determined by multiple unpaired t tests.

Data represent independent experiments with cells isolated from 3 donors.

All data represent the mean ± SD.

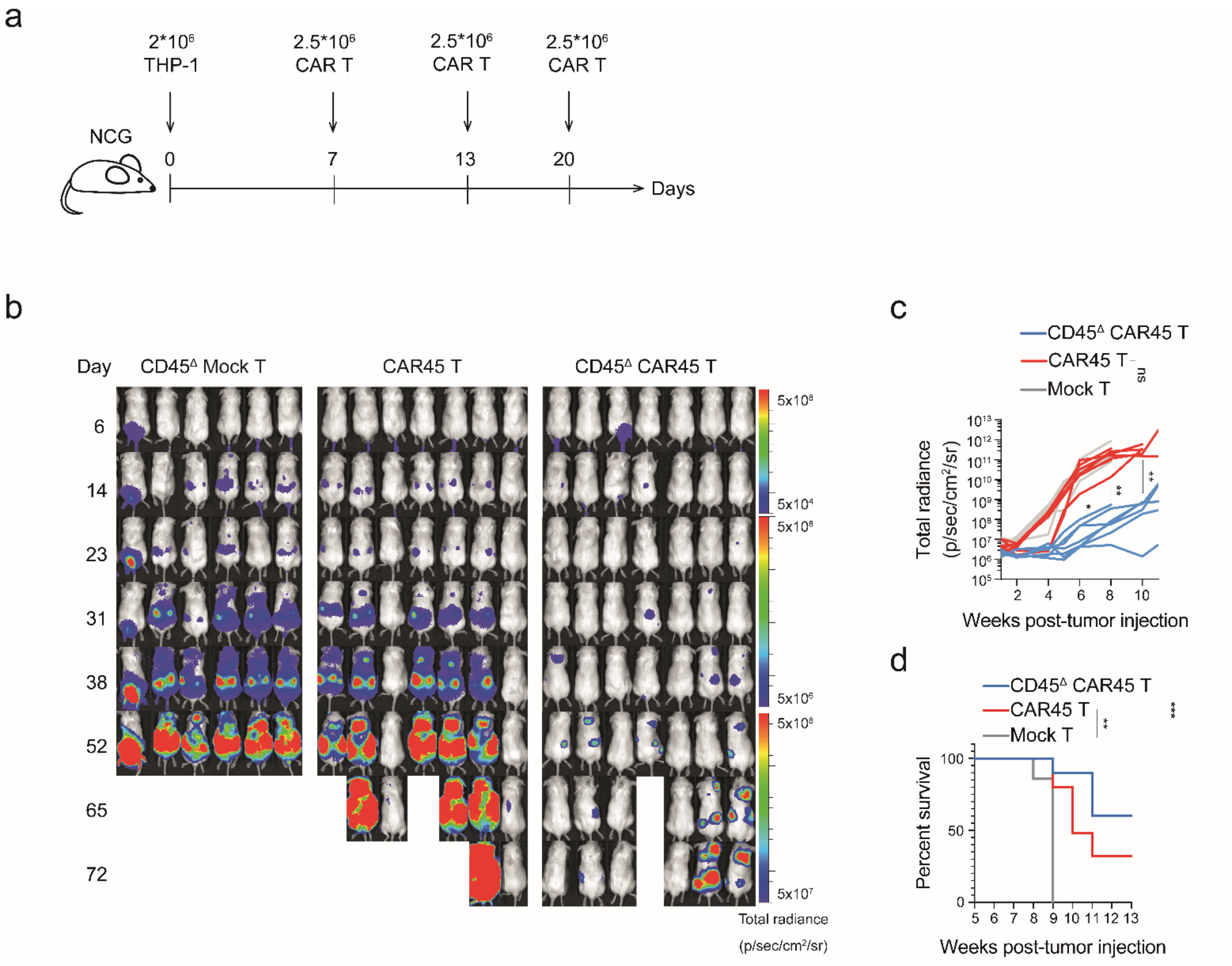


**Supplementary figure 3. CD45$^{\Delta}$ CAR45 T cells outperform CAR45 T cells *in vitro* and *in vivo*.**

**a.** NCG mice were subjected to i.v. infusion of 2x10$^{6}$ THP-1/ffluc cells. On days 7, 13 and 20 after tumor inoculation, animals were subjected to *i.v.* infusion of 1.5x10$^{6}$ mock, CAR45 or CD45$^{\Delta}$ CAR45 T cells.

**b.** Representative IVIS images of the mice from the mock, CAR45 and CD45$^{\Delta}$ CAR45 T-cell-treated groups.

**c.** Quantification of tumor burden (as the total radiance due to luciferase activity per mouse) from **b** for the 13-week period. P values were determined by multiple unpaired t tests. Nonsignificant values are not shown.

**d.** Kaplan–Meier curve showing overall animal survival in each experimental group. P values were determined by the log-rank Mantel–Cox test.

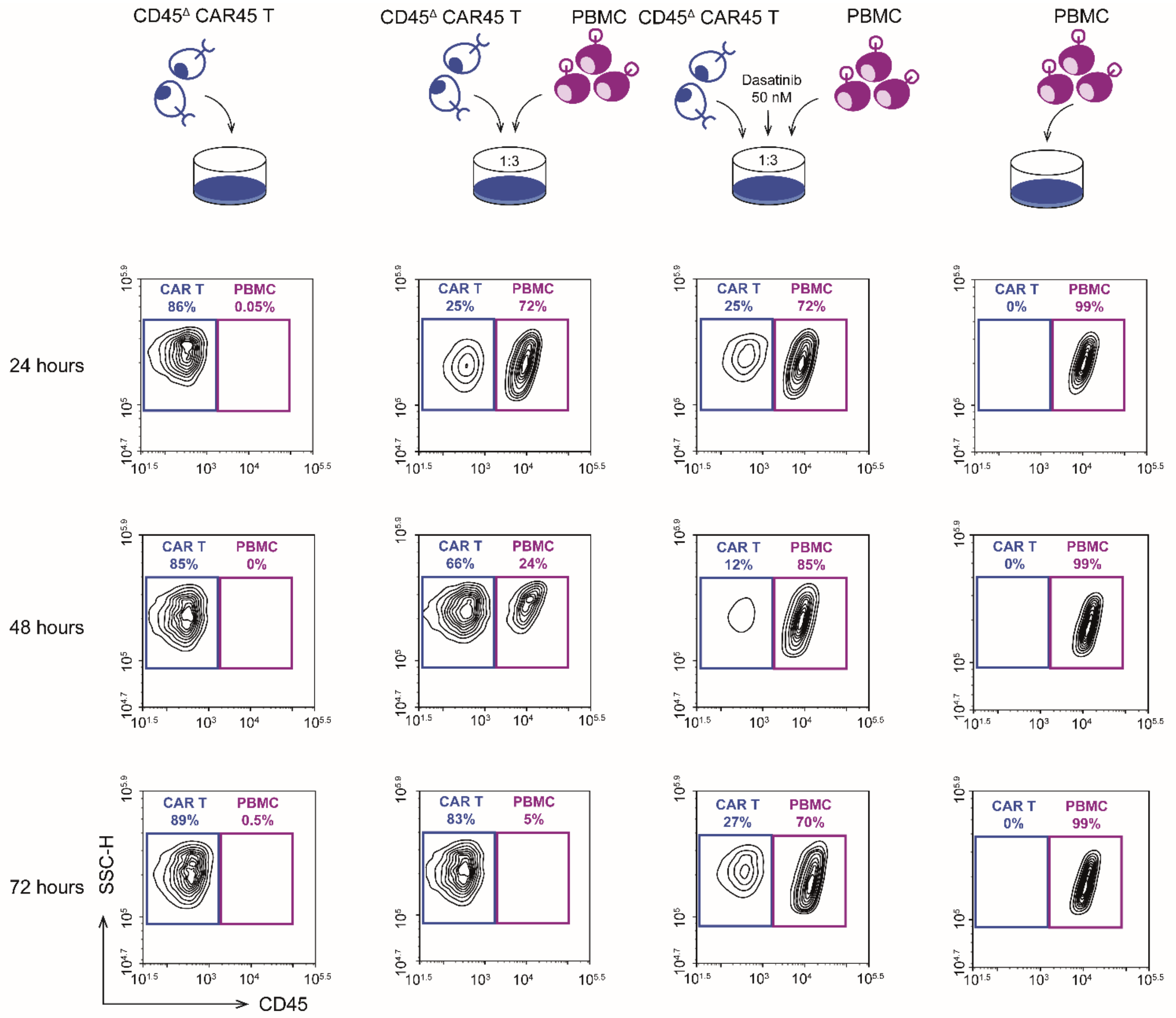


**Supplementary figure 4. Cytotoxicity assay of CD45$^{\Delta}$ CAR45 T cells against human PBMCs.** Freshly isolated PBMCs were cocultured with CD45$^{\Delta}$ CAR45 T cells at a 1:3 ratio for 24, 48 and 72 hours upon the addition of 50 nM dasatinib or no drug. Contour plots show the frequency of live PBMCs at the end of the coculture period. Control groups represent PBMCs or CD45$^{\Delta}$ CAR45 T cells cultured separately.

Data represent independent experiments with cells isolated from 3 donors.

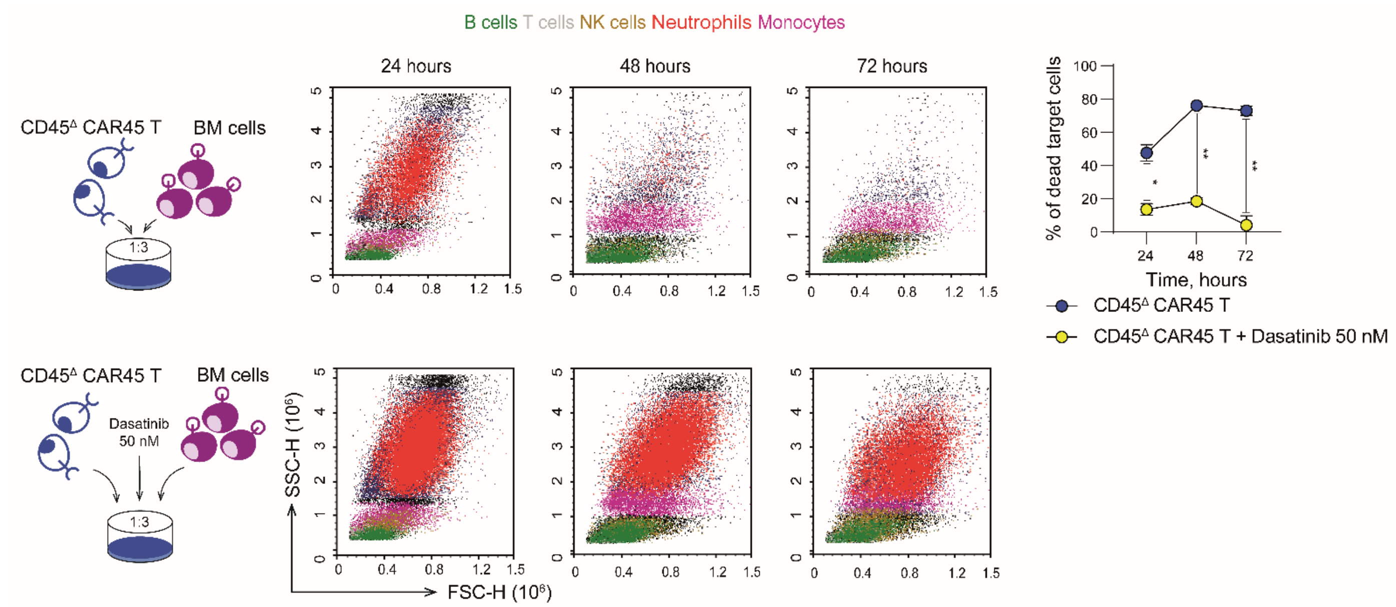


**Supplementary figure 5. Cytotoxicity assay of CD45$^{\Delta}$ CAR45 T cells against human BM cells.** Freshly isolated BM cells were cocultured with CD45$^{\Delta}$ CAR45 T cells at a 1:3 ratio for 24, 48 and 72 hours upon addition of 50 nM dasatinib or no drug. Contour plots show different populations of live BM cells at the end of the coculture period. Total numbers of live target cells were quantified by flow cytometry at 24, 48 and 72 hours. P values were determined by multiple unpaired t tests.

**Supplementary Table 1. Antibodies.**

| Antibody | Source | Identifier |
|---|---|---|
| anti-human CD45-APC-Cy7 | Sony | Cat.# 2120070 |
| anti-human CD45-PerCP | BD Biosciences | Cat.# 345809 |
| anti-human CD45-BV786 | BD Biosciences | Cat.# 563716 |
| anti-human CD45-PE-Vio 615 | Miltenyi | Cat.# 130-110-777 |
| anti-human CD4-FITC | Biolegend | Cat.# 317416 |
| anti-human CD4-APC-Cy7 | Thermo Fisher Scientific | Cat.# A15441 |
| anti-human CD4-APC-A750 | Beckman Coulter | Cat.# A94682 |
| anti-human CD4-KO525 | Beckman Coulter | Cat.# A96417 |
| anti-human CD8a-PE | Biolegend | Cat.# 300908 |
| anti-human CD8a-PE-Cy7 | Beckman Coulter | Cat.# 6607102 |
| anti-human CD45 | Abcam | Cat.# ab214437 |
| anti-human GAPDH | Abcam | Cat.# ab185059 |
| anti-human CD137-APC | Biolegend | Cat.# 309810 |
| anti-human IL2-APC-Cy7 | Biolegend | Cat.# 500342 |
| anti-human IFNγ-PE-Cy7 | Biolegend | Cat.# 502528 |
| anti-human CD45RA-PE | Biolegend | Cat.# 304108 |
| anti-human CD62L-APC | Biolegend | Cat.# 304810 |
| anti-human CD62L-BV711 | BD Biosciences | Cat.# 740783 |
| anti-human CD44-APC | Biolegend | Cat.# 397506 |
| anti-human CD3-APC | Biolegend | Cat.# 300312 |
| anti-human CD3-FITC | Biolegend | Cat.# 317306 |
| anti-human CD3-KO525 | Beckman Coulter | Cat.# B00068 |
| anti-human CD3-VioGreen | Miltenyi | Cat.# 130-113-142 |
| anti-human TIGIT-APC | Biolegend | Cat.# 372706 |
| anti-human TIM3-PE | Biolegend | Cat.# 345006 |
| anti-human LAG3-FITC | Biolegend | Cat.# 369308 |
| anti-human CD56-BV510 | BD Biosciences | Cat.# 563041 |
| anti-human IgG Fc Cross-Adsorbed Secondary Antibody, DyLight 650 | Invitrogen | Cat.# SA5-10137 |
| CD19 CAR Detection Reagent, human | Miltenyi | Cat.# 130-129-550 |
| anti-mouse CD45-PE | Sony | Cat.# 1115530 |
| Goat Anti-Rabbit IgG Antibody, HRP-conjugate | Sigma | Cat.# A0545-1ML |

**Supplementary Table 2. Off-target sites of nuclease activity with gRNA-2.**

| Target gene | Position | Part | CFD Score | Nuclease activity | # | Reads with indels, percents | | | | | |
|---|---|---|---|---|---|---|---|---|---|---|---|
| | | | | | | knockout + gRNA-2 | | | knockout - gRNA-2 | | |
| | | | | | | 4h | 8h | 16h | 4h | 8h | 16h |
| PTPRC | chr1:198639272 | exon | 1.00 | on-target | - | ***73,70*** | ***83,80*** | ***90,60*** | 1,50 | 1,70 | 1,60 |
| GPATCH4 | chr1:156594924 | exon | 0.50 | | #1 | 1,20 | 1,10 | 1,20 | 1,10 | 1,20 | 1,20 |
| CHST10\|NMS | chr2:100451099 | intergenic | 0.27 | | #2 | 1,30 | 1,30 | 1,30 | 1,30 | 1,20 | 1,30 |
| SNORD18\|AC096559.1 | chr2:12066072 | intergenic | 0.22 | | #3 | 0,70 | 0,70 | 0,70 | 0,60 | 0,60 | 0,70 |
| AC107057.1\|LINC01248 | chr2:5579273 | intergenic | 0.30 | | #4 | 1,70 | 1,90 | 1,80 | 1,80 | 1,80 | 1,90 |
| OSBPL6 | chr2:178393480 | intron | 0.22 | | #5 | 0,90 | 0,80 | 0,80 | 0,90 | 1,00 | 0,90 |
| NFKBIZ | chr3:101846553 | intron | 0.29 | | #6 | 1,20 | 1,20 | 1,20 | 1,10 | 1,10 | 1,20 |
| ATR | chr3:142500133 | intron | 0.29 | | #7 | 2,80 | 2,90 | 2,80 | 2,90 | 2,70 | 2,80 |
| TADA2B | chr4:7053277 | exon | 0.15 | | #8 | 0,80 | 0,70 | 0,80 | 0,80 | 0,70 | 0,70 |
| CTD-3179P9.2\|CTD-2281M20.1 | chr5:118371920 | intergenic | 0.21 | | #9 | 1,20 | 1,10 | 1,20 | 1,00 | 1,00 | 1,20 |
| SLC25A48 | chr5:135872223 | exon | 0.01 | | #10 | 0,50 | 0,50 | 0,50 | 0,50 | 0,50 | 0,50 |
| AL391416.1\|ME1 | chr6:83396465 | intergenic | 0.25 | | #11 | 1,80 | 1,90 | 1,90 | 1,90 | 1,90 | 1,80 |
| FBXL4 | chr6:98871971 | exon | 0.00 | off-target | #12 | 0,80 | 0,80 | 0,80 | 0,90 | 0,90 | 0,80 |
| AC000099.1 | chr7:127222790 | intron | 0.42 | | #13 | 0,30 | 0,40 | 0,30 | 0,20 | 0,30 | 0,30 |
| PLEKHA8 | chr7:30088697 | intron | 0.41 | | #14 | 0,70 | 0,70 | 0,90 | 0,60 | 0,70 | 0,70 |
| GPR141\|NME8 | chr7:37836988 | intergenic | 0.32 | | #15 | 0,40 | 0,40 | 0,50 | 0,40 | 0,50 | 0,40 |
| PVT1 | chr8:127986989 | intron | 0.22 | | #16 | 0,80 | 0,70 | 0,80 | 0,70 | 0,70 | 0,70 |
| PSD3\|AC100800.2 | chr8:18576436 | intergenic | 0.31 | | #17 | 1,20 | 1,10 | 1,20 | 1,10 | 1,00 | 1,00 |
| TMEM25 | chr11:118532157 | exon | 0.05 | | #18 | 1,30 | 1,30 | 1,30 | 1,40 | 1,40 | 1,40 |
| DCHS1/RP11-732A19.5\|DCHS1 | chr11:6624643 | intergenic | 0.24 | | #19 | 0,70 | 0,70 | 0,70 | 0,70 | 0,60 | 0,70 |
| SLC39A9 | chr14:69421114 | intron | 0.26 | | #20 | 3,00 | 3,90 | 3,70 | 3,50 | 2,70 | 3,20 |
| APBA2 | chr15:29051738 | intron | 0.25 | | #21 | 1,60 | 1,70 | 1,70 | 1,70 | 1,60 | 1,60 |
| THSD4\|THSD4/RP11-1123I8.1 | chr15:71504541 | intron | 0.69 | | #22 | ***1,40*** | ***1,50*** | ***1,70*** | 1,20 | 1,20 | 1,30 |
| HSPA13\|SAMSN1 | chr21:14430803 | intergenic | 0.50 | | #23 | 0,80 | 1,20 | 0,70 | 0,80 | 0,80 | 0,80 |
| CDKL5 | chrX:18433674 | intron | 0.47 | | #24 | 1,60 | 0,80 | 1,40 | 1,30 | 1,40 | 1,40 |

On-target activity is highlighted in light green. Detected off-target activity (by continuously increasing the percentage of reads with indels during the time after knockout with gRNA-2) is highlighted in light orange. The percentage of reads with indels for these cases is shown in ***italic bold***. For technical details of the analysis, please refer to the methods.
